# Spinal cord phosphoproteome of a SCA2/ALS13 mouse model reveals alteration of ATXN2-N-term SH3-actin interactome and of autophagy via WNK1-MYO6-OPTN-SQSTM1

**DOI:** 10.1101/2024.11.06.622233

**Authors:** Luis-Enrique Almaguer-Mederos, Arvind Reddy Kandi, Nesli-Ece Sen, Júlia Canet-Pons, Luca-Malena Berger, Jana Key, Melinda-Elaine Brunstein, Christian Münch, Suzana Gispert, Georg Auburger

## Abstract

Toxic polyglutamine (polyQ) expansions in ATXN2 trigger neurodegenerative processes, causing Spinocerebellar Ataxia type 2 (SCA2), and enhancing TDP-43-dependent pathology in Amyotrophic Lateral Sclerosis (ALS) / Fronto-Temporal Dementia (FTD). Primary disease events can be compensated transiently, delaying disease manifestation. To define potential therapy targets, we documented how cells modify their phospho-signals and how the ATXN2 interactome changes, using preferentially affected nervous tissues from end-stage *Atxn2*-CAG100-KnockIn mice. The spinal cord phosphorylome revealed massive hyperphosphorylations flanking the polyQ expansion in ATXN2 and for SQSTM1, and moderate hyperphosphorylations also for ALS proteins OPTN, UBQLN2, TNIP1 and TBK1-targeted TAX1BP1, versus strong hypophosphorylations of WNK1, SPARCL1 and PSMD9. Significant enrichments of SH3-containing proteins, autophagy / endocytosis factors, and actin modulators could be explained by N-terminal, polyQ-adjacent, proline-rich motifs in ATXN2. Coimmunoprecipitation profiling in cerebellum documented known associations with RNA-binding proteins like PABPC1 and TDP-43 with its modifier PPIA to decrease upon expansion, contrasting with increased binding of SH3-proteins, like MYO6, RPL21 and DLG4. Validation of protein and mRNA levels in mouse spinal cord, and embryonic fibroblasts or patient fibroblasts after bafilomycin or arsenite treatment, observed polyQ-dependent OPTN deficiency and SQSTM1 induction impairment. Overall, this combined phosphoproteome / interactome study efficiently revealed key pathways and molecular events.

## 1. Introduction

Spinocerebellar ataxia type 2 (SCA2) is a rare neurodegenerative disorder with autosomal dominant inheritance, caused by a (CAG)n repeat expansion mutation in the Ataxin-2 gene (human gene symbol *ATXN2,* chromosomal cytogenetic band 12q24.12), encoding an expanded polyglutamine (polyQ) tract near the N-terminus of Ataxin-2 protein [1–3]. Repeat expansions beyond (CAG)32 repeats lead to typical spinocerebellar manifestation, by affecting glutamatergic synapses preferentially. This occurs in two main areas: in the spinal cord, where it impacts connections between upper and lower motor neurons, and in the cerebellum, where it disrupts the synapses between granule neuron parallel fibers and the dendritic trees of Purkinje neurons. Subsequent atrophy of further neuronal circuits, denervation of skeletal muscles and demyelination, with final patient immobility, leads to a characteristic combination of tremor, leg cramps, slowed saccadic eye movements, decreasing body mass index, and death within 30 years after clinical manifestation [4–8]. While unable to cause the monogenic, fully penetrant SCA2, the intermediate-length expansions with size (CAG)27-32 act within polygenic networks to increase the disease risk and/or progression of (i) amyotrophic lateral sclerosis (ALS or ALS13) [9–14], (ii) frontotemporal dementia (FTD) [15–17], (iii) tauopathies named as progressive supranuclear palsy (PSP) or as Parkinson plus [18], and also (iv) idiopathic Levodopa-responsive Parkinson’s disease [19]. ALS and FTD display a characteristic neuropathological combination of cortical and spinal motor neuron degeneration in the absence of cerebellar atrophy, with the appearance of cytosolic inclusion bodies that contain the normally nuclear ribonucleoprotein TDP-43 as hyperphosphorylated, ubiquitinated, and proteolytically cleaved C-terminal fragment [20, 21]. This ribonucleoprotein/RNA aggregation process in ALS/FTD usually occurs without co-localization of expanded ATXN2, except transiently in individual patients [22–25]. In view of the causal neurotoxic role of ATXN2 for SCA2, and of its modifier role for ALS and FTD, the most promising approaches towards their preventive neuroprotective treatment are focused on repressing the expression of *ATXN2* mRNA. Successful pre-clinical studies showed the KnockOut (KO) of murine Ataxin-2 to extend the survival of TDP-43 overexpressing mice from 20 days to more than 2 years [26]. The *in vivo* administration of antisense oligonucleotides (ASO) or Cas13 CRISPR effectors to eliminate murine *Atxn2* mRNA was found to improve functional deficits, extend survival, and reduce neuropathology in ALS models [26–28]. Similarly, pre-clinical studies conducted in SCA2 mouse models also showed ASO intrathecal delivery as a promising disease-modifying treatment for this condition [29]. Furthermore, the identification of small drugs that minimize Ataxin-2 expression was beneficial in a SCA2 mouse model, by normalizing molecular markers of pathology such as the levels of the selective autophagy receptor (SAR) SQSTM1, also known as p62 [30]. Conversely, the lentiviral introduction of expanded ATXN2 into brain triggers the accumulation of LC3-II and SQSTM1 as aggrephagy mediators [31, 32], while pharmacological autophagy enhancement is protective in SCA2 models [33]. Thus, clinical trials to prevent SCA2 and ALS via Ataxin-2 knockdown approaches have been initiated, and will need to be optimized by the identification of the neurotoxic Ataxin-2 interactions, and by the definition of molecular events that serve as biomarkers for progression versus regeneration.

What are the binding partners and physiological roles of Ataxin-2, what is its connection with TDP-43, and how does ATXN2 acquire neurotoxicity upon polyQ expansion? It is relevant to know that the KO of Ataxin-2 results only in age-associated obesity, hypercholesterolemia, and hepatosteatosis [34–36], whereas the KO of its more abundantly expressed paralog Ataxin-2-like (ATXN2L) causes mid-embryonic lethality [37]. Vertebrates (except birds) have developed two independent Ataxin-2 gene copies, while only one gene copy exists from nematode worms (*ATX-2*) via fruit flies (d*Atx2*) to yeast (*PBP1*), and plants like *Arabidopsis thaliana* again have two copies (CID3 and CID4) [38]. This strong sequence conservation across eukaryotes is most evident in several functional domains: (i) As a common element in all Ataxin-2 family members, the poly(A)-binding protein associated motif 2 (PAM2) and its interaction with the MLLE domain of poly(A)-binding protein 1 (PABPC1) was first identified in yeast. Using this knowledge, various studies convincingly demonstrated the impact of the Ataxin-2 family on mRNA poly(A) tail length, translation efficiency, and ribonucleoprotein association [39–53]. (ii) All Ataxin-2 family members have a domain with similarity to the Sm motif of nuclear splice factors, although Ataxin-2 and several other proteins with such an LSm (like Sm) domain localize in the cytosol where no splicing occurs [54–56]. It was shown that this LSm domain and the LSm-associated domain (LSm-AD) in Ataxin-2 mediate direct binding and processing of RNAs, and interaction with RNA processing proteins such as the RNA helicase DDX6 [57–60]. Together, the PAM2 motif and the LSm domain explain why the Ataxin-2 family members relocalize with PABPC1 / mRNAs / ribonucleoproteins (e.g. TIA1, G3BP1) / 40S ribosome subunits to condensate in stress granules, when cellular damage such as oxidative stress or starvation requires the quality control and possibly degradation of RNAs [61–63]. The indirect association of Ataxin-2 with the ALS disease protein TDP-43 is thought to be mediated by RNAs that are associated with the LSm / LSmAD domains [9, 28, 64]. (iii) The presence of a long polyQ domain within Ataxin-2 is prominent in primates and insects, but shorter polyQ domains appear in most mammalian ATXN2 orthologs, at differing positions [38]. (iv) Proline-rich motifs (PRM) are frequent in several positions among the Ataxin-2 family members [38]. At least four distinct PRM exist within human Ataxin-2, two within the main isoform after the polyQ repeat, and two or three before the polyQ repeat within the extended N-terminus of an alternative longer isoform, usually having the classical sequence **P**XX**P**XX<u>R</u> or <u>R</u>XX**P**XX**P** that can interact with SRC-homology domain type 3 (SH3) domains. Their SH3-containing interactor proteins control actin filament assembly itself, or the interaction of actin fibers with the plasma membrane receptor tyrosine kinase (RTK) endocytosis apparatus. Indeed, mostly the third PRM in ATXN2 was demonstrated to associate with the SH3-containing proteins CIN85, ITSN1, SRC, and Endophilin A1/A3. Via this association, Ataxin-2 repressed the internalization of the Epidermal Growth Factor Receptor (EGFR) in peripheral cells, and would be expected to have analogous effects on other RTK family members in the nervous system [65–68], so that ATXN2 orthologs from yeast via nematodes to mammals were shown to repress mTORC1 growth signals and cell size, modulating autophagy via ULK1 [48, 69–74]. Overall, the principal phenotypes of d*Atx2* mutation in fruit flies are (i) impaired translation due to altered direct association of Ataxin-2 with polyribosomes, and (ii) aberrant sensory bristle morphology due to an indirect and poorly characterized influence of Ataxin-2 on actin filament formation [42, 75]. Until the present study, it had remained unclear to what degree these two functions are critical for the disease, and how they are affected by the initial partial lossof-function or the subsequent neurotoxic and progressive gain-of-function of polyQ expansion disorders.

To model SCA2 and ALS13 disease processes upon ATXN2 polyQ expansion by a genetic approach in animals, we used targeted recombination to perform the KnockIn (KIN) of a (CAG)100 repeat at the appropriate position of the *Atxn2* gene within the mouse genome. An additional insertion of LoxP sites was performed at either side of the (CAG)100 repeat, which permits the conditional conversion of the KIN-triggered neurodegeneration into a KO that would halt the neurotoxicity and permit partial regeneration, once a transgenic Cre recombinase in such mouse mutants is activated by Tamoxifen injection, at any disease stage [76]. This approach was successful in generating the *Atxn2*-CAG100-KIN mouse mutant where the physiological promoter governs endogenous Ataxin-2 expression regulation, and where the expansion is unstable over generations, just as in patients. Faithfully mirroring the temporal and spatial pattern of SCA2 pathology, the polyQ expansion in ATXN2 is detectable in the central nervous system and in peripheral liver tissue of these mutant mice [77–80], triggering initial obesity by the age of 10 weeks (reflecting an initial partial loss-of-function) that develops into weight loss by 10 months (reflecting the subsequent progressive gain-of-function), with clear deficits of vertical movement coordination by the cerebellum from 4 months, and a reduction of grip strength as a reflection of motor neuron degeneration from 9 months onward [77, 81]. In the spinal cord, progressive TDP-43 aggregation is observed, together with cholesterol anomalies [81]. In the cerebellum, prominent anomalies of calcium and sphingomyelin lipids are documented, as in SCA2 patients [79, 80]. Overall, the breeding of homozygous mouse mutants causes a more severe progression than heterozygotes, similar to the unusually strong affection of homozygous SCA2 patients [77, 82–84]. The lifespan of the homozygous KIN mice is limited to 14 months, while matched wildtype (WT) mice have a life expectancy well beyond 24 months [77]. In contrast to SCA2 mouse models with transgenic overexpression of recombinant ATXN2, this KIN mouse can be expected to reflect the anomalies of Ataxin-2 splicing, proteolytic cleavage, post-translational modifications, and interactor binding, in patterns that would be characteristic for the human SCA2 and ALS13 nervous system. Such investigations in patient tissues are usually impossible, due to the unavailability of high-quality autopsy material in such rare disorders.

In the present study, a survey of phosphorylation changes was documented in the spinal cord of such KIN mice at the terminal disease stage. As technical approach, the phosphorylated peptides from a proteolytically cleaved protein extract were bound by iron-nitrilotriacetic acid (Fe-NTA) on immobilized metal affinity chromatography (IMAC) columns, to be subsequently sequenced and quantified by label-free mass spectrometry. We hoped to elucidate how the tyrosine (Tyr, Y) phosphorylations at the receptor endocytosis machinery are adapted, and how the cells adapt their stress signaling via serine (Ser, S) / threonine (Thr, T) phosphorylation cascades from the plasma membrane to the nucleus, attempting to counteract the specific atrophy process of SCA2 at crucial molecules and pathways. To distinguish primary pathological events within the ATXN2 interactome from secondary downstream consequences and compensatory efforts, ATXN2 coimmunoprecipitates were sequenced with quantitative comparison between *Atxn2*-KO, *Atxn2*-WT, and *Atxn2-CAG100-KIN* cerebellum.

## 2. Results

### 2.1. Phosphoproteome profile of end-stage spinal cord from authentic SCA2/ALS13 mouse model

Analyzing biological replicates of four WT versus four *Atxn2*-CAG100-KIN spinal cord tissues from 14-month-old mice, Fe-IMAC phosphorylation profiling detected a total of 80,278 redundant modified peptide assignaments to 27,105 modified sites, and documented 428 significantly downregulated versus 552 significantly upregulated phospho-peptides with at least 2-fold changes (Table S1A-E).

Upon their analysis with the STRING webserver (Table S1F), enrichments of pathways and Gene Ontology terms were observed, with significances among the hyperphosphorylations for autophagy, ubiquitin, proteasome, ALS, and synapses (scheme of autophagy phospho-regulation in Fig. 1A and Table S1G), while additional enrichments among hypophosphorylations for ion channels, glutamate neurotransmission, and myelin factors presumably reflected selective neuron loss.

**Figure 1:**
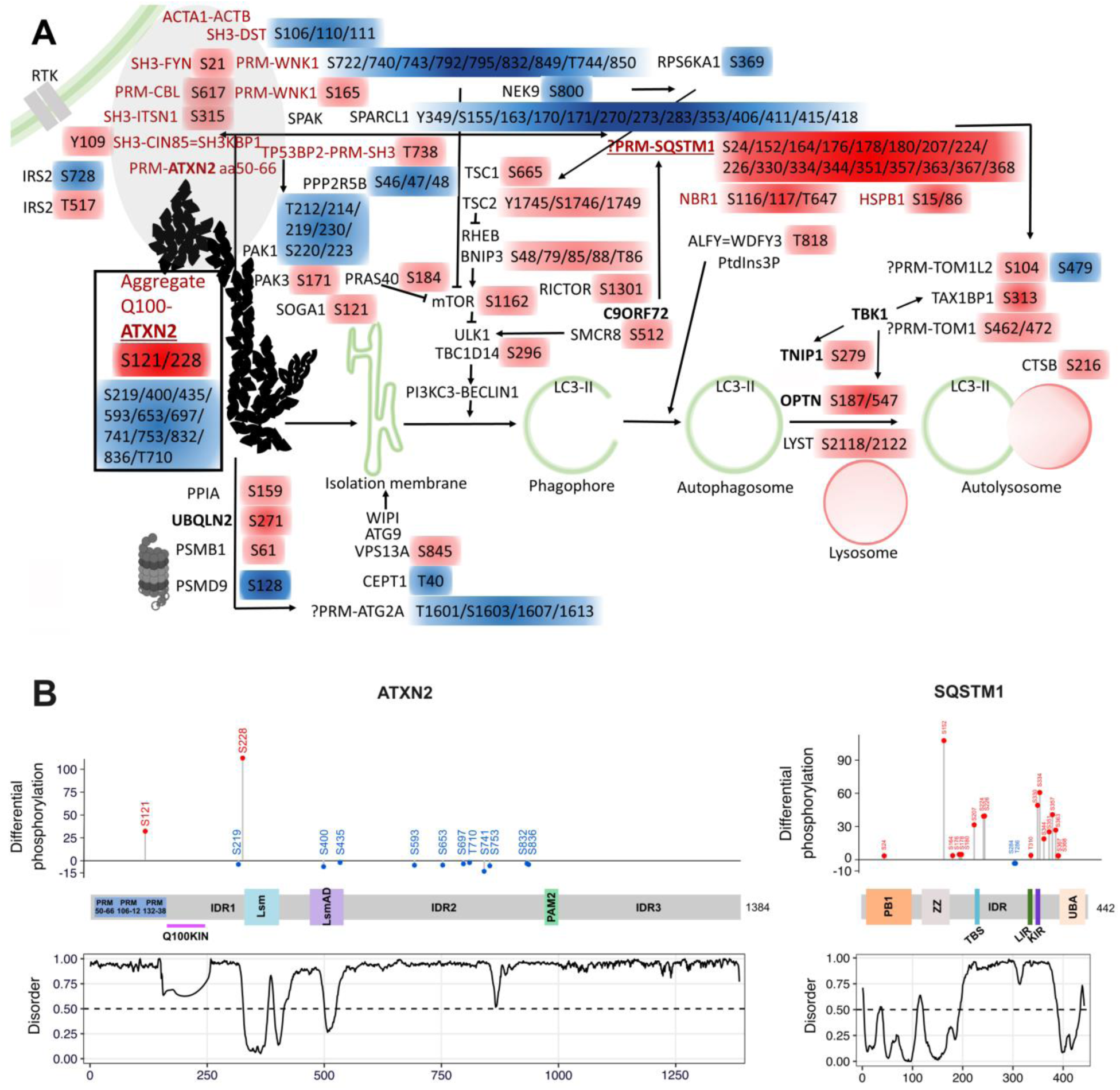
**(A)** Schematic summary of phospho-regulation events of proteolysis and aggrephagy in the terminal-stage spinal cord of the *Atxn2*-CAG100-KIN mouse model of SCA2/ALS13. The potential co-aggregation of ATXN2 (black clusters) with SH3-containing interactors and competing PRM-containing proteins that show differential phosphorylation is highlighted in a field with a gray background. Proteins potentially undergoing LLPS are highlighted in red letters, ALS disease proteins in bold letters. **(B)** Schematic protein structures and Q100-ATXN2-triggered differential phosphorylation events for ATXN2 as aggregating disease protein and for SQSTM1 as the main selective aggrephagy receptor, illustrating that massive hyperphosphorylations are flanking the Q100-KIN mutation replacing murine residue Q156 (purple line) and the three PRMs (deep blue square, with mouse residue numbers) in the N-terminus of ATXN2, and are targeting the ZZ (light gray square) and LIR (deep green column) domains of SQSTM1. Hypophosphorylations across remaining ATXN2 likely reflect its reduced abundance upon polyQ expansion [77]. The numbering of residues adheres to UniProt reference sequences, ignoring the unstable polyQ repeat KIN in ATXN2. Proteins are represented by their gene symbols, phosphorylated residue numbers are shown for serine (S), threonine (T) or tyrosine (Y). IDR = intrinsically disordered region, prone to phase separation; LIR = LC3-interacting region; KIR = Keap1-interacting region; PB1 = Phox and Bem1p; PtdIns3P = phosphatidylinositol 3-phosphate; RTK = receptor tyrosine kinase; TBS = TRAF6-binding sequence; UBA = ubiquitin-associated; ZZ = Zinc finger domain.

As the key result, the ATXN2 N-terminus with its polyQ expansion and three presumptive PRMs (UniProt O70305 residues 50-66, 106-12, 132-38) exhibited two massive hyperphosphorylations (112-fold at Ser228, 32-fold at Ser121). In phosphorylome mass spectrometry databases, we found no previous reports of phosphorylation or hyperphosphosphorylation at these residues, so they differ from known signaling cascades. Thus, they may represent efforts to prevent phase separation and reflect the progressive neurotoxic ATXN2 gain-of-function in SCA2/ALS13. Similarly massive hyperphosphorylations were observed only in the Zinc finger domain (ZZ) and LC3-interacting region (LIR) domains of aggrephagy receptor SQSTM1 (Fig. 1B). Conversely, eleven hypophosphorylations (at least -2-fold) across ATXN2 from LSm domain to C-terminus (Fig. 1B) correspond to the known reduced abundance of Q100-ATXN2 in KIN nervous tissue due to impaired transcription/translation [77]. They may reflect the initial partial ATXN2 loss-of-function phenotypes in SCA2.

Among all hyperphosphorylation events, enrichment of SMART protein motifs by STRING analysis revealed SRC homology type 3 (SH3) domains to be overrepresented with a false discovery rate (FDR) of 4.09e^−08^, including hyperphosphorylation of known ATXN2 interactors such as CIN85 and ITSN1 [65, 67, 68]. This may be caused by pathogenic interactions of ATXN2 PRMs with SH3 domains in the RTK / actin pathways, rather than its intrinsically disordered regions (IDR) [85], which trigger LLPS.

### 2.2. Detailed analysis of pathway enrichments for SH3 domains and for autophagy factors

As listed in Table S1I-L, differential phosphorylation was firstly documented for 28 SH3-domain-containing proteins (e.g. hyperphosphorylations at a Tyr residue for ATXN2-interactor CIN85 aka SH3KBP1, and at different Ser residues for CIN85 and ITSN1 as endosomal regulators, for TP53BP2 as autophagy modulator, for SHANK1 and the tyrosine kinase FYN as regulators of glutamatergic neurotransmission and myelination, versus hypophosphorylations for DST as autophagosome transport factor).

Differential phosphorylation was secondly noted also for 40 proteins with functionally studied PRM (like some hyper- and many strong hypo-phosphorylations of WNK1 as autophagy modulator), and for 50 proteins with candidate PRM sequences near their dysregulated phospho-sites (Table S1K, including the hyperphosphorylations in SQSTM1), which have the potential to enter in aberrant competition with PRMs of ATXN2. This is very relevant for aggrephagy because the interactions of PRMs with SH3 domains are a classical driver of LLPS that promotes protein aggregation [86–92].

We attempted to elucidate, which sequences in ATXN2 conform with the 10 established subtypes of PRMs (Fig. 1 in [93]), being potentially responsible for aberrant binding to SH3-containing proteins and for abnormal competition with other PRM-containing proteins, which exhibit differential phosphorylation. Filtering all phosphoproteome data for experimentally confirmed PRM-SH3 interactions, the most N-terminal ATXN2 PRM (murine amino acids 50-66, human aa 55-71) fitted the sequence constraints to explain interaction with almost all SH3-domains with differential phos-phorylation, and another N-terminal ATXN2 PRM (murine aa 132-138, human aa 142-148) explained the NCF2-CYBA interaction (Figure 2). Thus, differential phosphorylation in direct interactor chains via PRM-SH3-PRM bridges may reflect the selective impact of Q100-ATXN2 on glutamatergic postsynapses (SHANK1, GRM1), myelination (FYN, MBP) and autophagy (WNK1).

**Figure 2:**
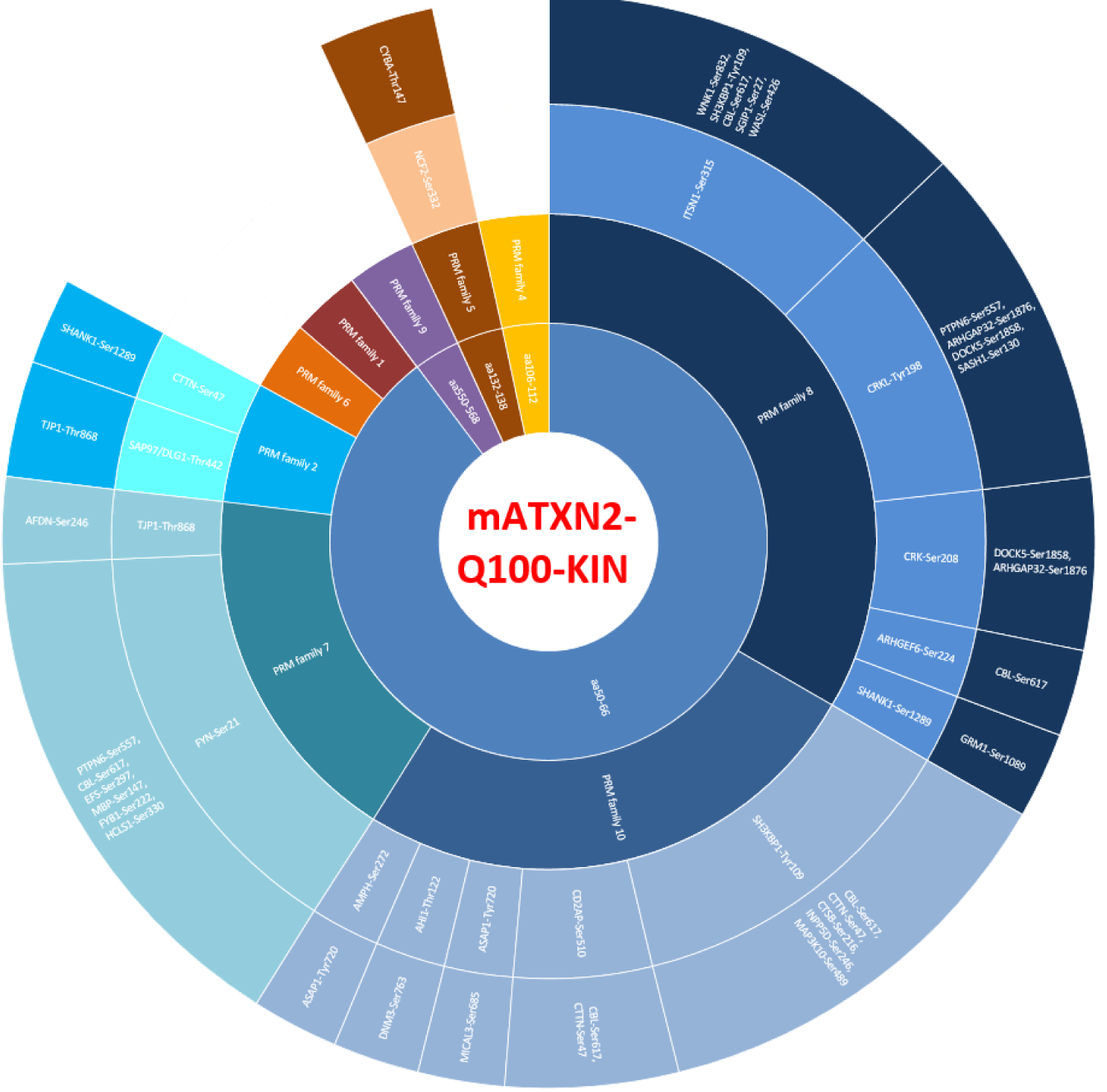
Synopsis of differentially phosphorylated PRM-containing and SH3-containing proteins, in their experimentally verified interactions according to current literature, and in their binding to subtypes of PRM sequences within human ATXN2. In this sunburst diagram, the outer ring shows PRM-containing differentially phosphorylated proteins that may compete with ATXN2 for SH3 binding. At appropriate positions in the ring below, their established SH3-containing interactors are shown with their phosphorylation changes. In the middle ring, the respective family of PRM sequences is shown that would bind to each SH3-containing protein. Inner rings illustrate which N-terminal ATXN2 sequences correspond to these PRM families, and could therefore compete with the PRM-containing proteins in the outer ring for their SH3-associations. Within ATXN2, the PRM at murine residues 50-66 has a complex pattern of prolines that can interact with diverse SH3-domains, such as PRM-subtypes 2 (may impair e.g. CTTN binding to SHANK1), 7 (may impair FYN binding to FYB1 and MBP and PTPN6), 8 (may impair CIN85/SH3KBP1 binding to CBL) and 10 (may impair SHANK1 binding to GRM1, and also ITSN1 binding to WNK1).

The 7.2-fold hyperphosphorylation of tyrosine kinase FYN (whose SH3 motifs would interact with ATXN2 proline-rich motifs (PRM) similar to SRC as its family member [65, 67]) can influence endosome dynamics, nutrient uptake and axon myelination (with strongly hyperphosphorylated BCAS1 and hypophosphorylated MBP / MAG / MOBP), conspiring with hyperphosphorylations of TSC2 to control mTORC1 signals, autophagy initiation and cell growth [94, 95]. This is the same pathway where ULK1 and ALS protein C9ORF72 were found to synergize with ATXN2 [96–98].

Differential phosphorylation was thirdly encountered at tyrosine residues in 26 proteins, and indeed the tyrosine phosphatase PTPN6 showed a 2.1-fold pSer557 change. These observations suggest trophic signaling at receptor tyrosine kinases to be altered (like 2.6-fold pTyr1745-TSC2 as mTOR repressor, 4.0-fold pTyr3363-ANK2 as actin adaptor in motor neurons, or -4.6-fold pTyr47-NDRG2 as neuronal differentiation factor) (Table S1I-L).

As a final important result of STRING enrichment statistics, the PB1 domains that mediate multimerization of selective autophagy receptors were overrepresented (FDR 0.0159).

Among the top 30 hyperphosphorylations (>7-fold), the ALS disease proteins ATXN2, SQSTM1, UBQLN2, OPTN, and TBK1-regulated TAX1BP1 stood out (compared to the ALS protein TNIP1 with 2.7-fold hyperphosphorylation, see Table 1 and Table S1H). This was in overlap with 4 selective autophagy receptors that mediate [99, 100] aggrephagy, lysophagy, pexophagy and granulophagy (SQSTM1, NBR1 and TAX1BP1; further, WDFY3 showed a 3.2-fold increase, Table S1E) as well as mitophagy (OPTN; with four peptides from BNIP3 showing 2.4 to 2.2-fold increases). Further proteostasis factors among the top 30 hyperphosphorylations included the proteasomal subunit PSMB1, the lysosomal peptidase CTSB, and the LC3-dependent phagocytosis factor CYBA [101, 102] (Table 1). Particularly strong reductions of phosphorylation were documented in WNK1 (fold change -7.1) and in NEK9 (fold change -3.8), an adaptor for actomyosin autophagy in cooperation with MYH9 [103–105], suggesting cytoskeletal affection.

**Table 1.** List of the strongest hyperphosphorylations in KIN spinal cord (changes above 7-fold, showing aggregating disease protein in green letters, proteostasis factors in brown, actin-/myelin-associated factors in blue).

| fold change | Protein Symbol | Phospho- Sites | PubMed-ID<br>Autophagy | PubMed-ID<br>Actin | PubMed-ID<br>SCA-ALS |
| --- | --- | --- | --- | --- | --- |
| 112.2 | ATXN2 | Ser228 | 38308810,<br>30982600 | 12524342,<br>34977501 | 8896555,<br>28405022 |
| 107.6 | SQSTM1 | Ser152 | 22736434,<br>33332721 | 25015291 | 20301623,<br>35143965 |
| 66.2 | PSMB1 | Ser61 | 36602366 | - | - |
| 60.9 | CDR2L | Ser344 | - | 19427213,<br>19602412 | 36729270 |
| 60.7 | SQSTM1 | Ser330/Ser334/Ser344 | 31585693 | 25015291 | 30657305,<br>35143965 |
| 50.2 | ANK2 | Ser914 | 35226578 | 25533844,<br>29163198 | 29078305 |
| 49.1 | SQSTM1 | Ser330/Ser344 | 22736434,<br>33332721 | 25015291 | 30657305,<br>35143965 |
| 49.1 | SQSTM1 | Ser334/Ser344 | 22736434,<br>33332721 | 25015291 | 30657305,<br>35143965 |
| 47.4 | BCAS1 | Ser443 | - | 16133941,<br>29212715 | - |
| 40.7 | SQSTM1 | Ser357/Ser363/Ser368 | 22736434,<br>33332721 | 25015291 | 30657305,<br>35143965 |
| 39.3 | SQSTM1 | Ser226 | 22736434,<br>33332721 | 25015291 | 30657305,<br>35143965 |
| 39.3 | SQSTM1 | Ser224 | 22736434,<br>33332721 | 25015291 | 30657305,<br>35143965 |
| 39.2 | SQSTM1 | Ser357/Ser363/Ser367 | 22736434,<br>33332721 | 25015291 | 30657305,<br>35143965 |
| 32.2 | ATXN2 | Ser121 | 38308810,<br>30982600 | 12524342,<br>34977501 | 8896555,<br>28405022 |
| 31.5 | SQSTM1 | Ser207 | 22736434,<br>33332721 | 25015291 | 30657305,<br>35143965 |
| 26.7 | SQSTM1 | Ser363/Ser368 | 22736434,<br>33332721 | 25015291 | 30657305,<br>35143965 |
| 25.2 | SQSTM1 | Ser351/Ser363/Ser368 | 22736434,<br>33332721 | 25015291 | 30657305,<br>35143965 |
| 18.4 | SQSTM1 | Ser344 | 22736434,<br>33332721 | 25015291 | 30657305,<br>35143965 |
| 17.8 | PDHA1 | Ser295/Ser300 | 35731579 | - | 20002125,<br>37422902 |
| 17.2 | UBQLN2 | Ser271 | 35708843 | 23635659,<br>24549040 | 25398946,<br>30442662 |
| 16.6 | ARID1A | Ser1601 | 36229854 | 25195934,<br>38017409 | 28662958 |
| 14.6 | UBQLN2 | Ser267/Ser271 | 35708843 | 23635659,<br>24549040 | 25398946,<br>30442662 |
| 10.9 | CYBA | Thr147 | 33138004 | 16341922 | 23830905 |
| 10.7 | TAX1BP1 | Ser313 | 35470757,<br>33207181 | 29398621 | - |
| 10.0 | OPTN | Ser547 | 29867991,<br>33783320 | 23781020,<br>29875258 | 20428114,<br>38178841 |
| 9.8 | NBR1 | Ser117 | 21189453,<br>36255390 | - | 27551461,<br>31362587 |
| 8.7 | CTSB | Ser216 | 30107168 | 23018451 | 32757690 |
| 8.3 | SPP1 | Ser283/Ser290 | 21374592,<br>36505285,<br>38082480 | 10842093,<br>34576260 | 36460480 |
| 7.3 | CDH10 | Ser784 | 35721483 | 29030434 | 31176720 |
| 7.2 | FYN | Ser21 | 22745922 | 10224664,<br>28887403 | 28935570,<br>30154836 |

A unique massive hyperphosphorylation (107.6-fold) at Ser152-SQSTM1 was observed at the ZZ domain (Figure 1) as the cargo recognition motif for N-term-arginylated proteins, whose target association facilitates disulfide bond-linked multimerization and LC3-mediated autophagosome binding [106, 107]. This finding suggests an activation of arginyltransferase-1 (ATE1) to facilitate autophagic removal of polyQ-expanded ATXN2 via SQSTM1, as described for ER-phagy [108, 109]. Also actin assembly rates in linear and branched networks are modified by N-terminal arginylation [110, 111]. Indeed, ATE1 showed significant hyperphosphorylation in the present dataset.

In contrast, a striking paucity of ribonucleoproteins was apparent among all altered phosphosites (except ATXN2, with small changes for DCP1A, DDX3, EDC3, EDC4, PCBP1, PRKRA, RPS6KA1, SLIRP, SRRM1, SRSF6, SRSF9, TRIR), conspicuously showing no change for TDP-43, FUS, C9ORF72, TBK1, SOD1 or other ALS1-28 disease proteins. This may either suggest that ribonucleoprotein functions are not much affected, or that these factors are differently regulated, e.g. by arginine-methylation rather than by phosphorylation.

### 2.3. Levels of protein abundance and mRNA expression for dysregulated autophagy factors in endstage spinal cord from Atxn2-CAG100-KIN mice

The phospho-changes affected an individual residue e.g. in the case of ATXN2L, TAX1BP1, PPIA, CTSB, and WDFY3 (Fig. S1). However, ATXN2, SQSTM1, NBR1, as well as OPTN, SPARCL1, UBQLN2, BNIP3 and WNK1 showed similar dysregulations across several domains (Fig. 1 and S1). Therefore, it was important to assess (at least for key regulators) the overall protein abundance and transcript levels. Quantitative immunoblots of the KIN spinal cord tissue showed elevated abundance for SQSTM1, NBR1, BNIP3, WDFY3, CTSB, PPIA and TAX1BP1, but only the *Sqstm1* and *Ctsb* mRNA were increased upon RT-qPCR (Fig. 3 and S2/3). UBQLN2 abundance was unchanged (Fig. S3). There was no specific and sensitive antibody to detect endogenous WNK1 available to us. Despite the increase of its two phospho-sites, OPTN protein abundance was diminished, and its mRNA levels were unchanged (Fig. 3). Thus, these selective hyperphosphorylation events appear to reflect true cellular signaling efforts. This includes the 10.0-fold increase of pSer547, which is close to the C-terminal region (that binds MYO6 and TAX1 in competition with TAX1BP1 [112–114]) and its poly-ubiquitin binding ZF motif [115]. The meaningful OPTN hyperphosphorylations also include the 2.7-fold increase of pSer187 within the LIR domain, which regulates the binding to autophagosome membranes. Analysis of the MYO6 abundance in KIN spinal cord showed significantly elevated levels, but the *Myo6* mRNA was decreased (Fig. 3). This putative transcriptional compensation suggests the accumulation of autophagosome fusion factor MYO6 to be pathological. Consistent with its many decreased phospho-sites, SPARCL1 protein abundance and *Sparcl1* mRNA levels were reduced (Fig. 3). SPARCL1 is an extracellular factor that promotes excitatory synaptogenesis, as member of a protein family that controls autophagy and apoptosis [116, 117]. Transcript decreases were also observed for *Bnip3*, *Wdfy3, Ppia*, and *Tax1bp1*, suggesting that the encoded proteins have aberrantly slowed turnover, and that the accumulation of the encoded proteins has a pathological effect leading to compensatory transcriptional downregulation. Unchanged levels were documented for *Ubqln2* and *Wnk1* (Fig. S2).

**Figure 3.**
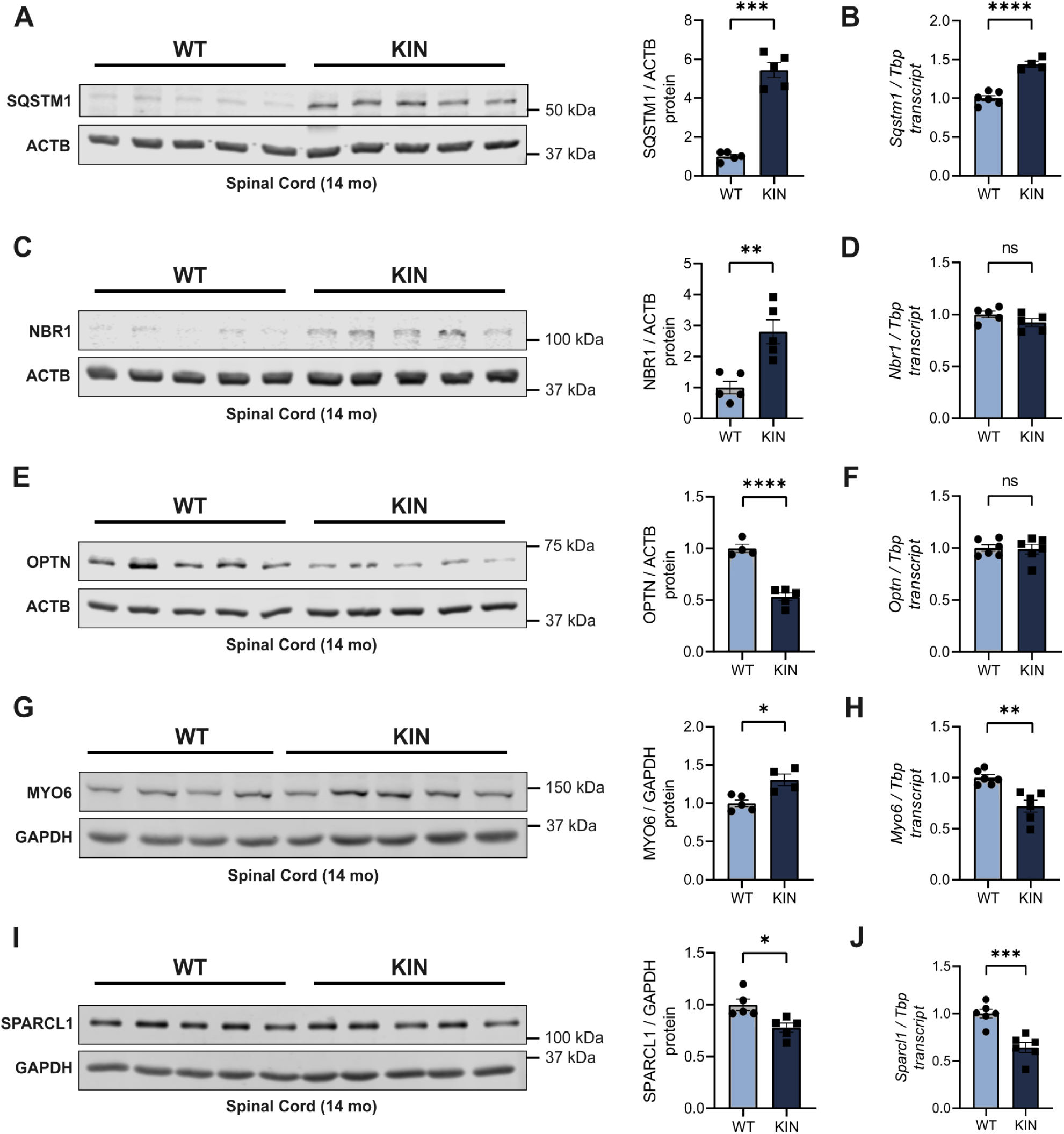
Quantitative immunoblots (**A**,**C**,**E**,**G**,**I**, N = 5 or 4 vs. 5) and quantitative reverse-transcriptase real-time PCR (**B**,**D**,**F**,**H**,**J,** N = 6 vs. 6) for SQSTM1 (**A**,**B**), NBR1 (**C**,**D**) and OPTN (**E**,**F**) as main aggrephagy receptors, as well as MYO6 **(G**,**H)** and SPARCL1 **(I**,**J)** in end-stage spinal cord from our SCA2/ALS13 mouse model *Atxn2*-CAG100-KnockIn (KIN) versus wildtype (WT). SQSTM1 and NBR1 were chosen in view of their hyperphosphorylated peptides, OPTN in view of its mix of hyper- and hypophosphorylations, together with its interactor MYO6, and SPARCL1 in view of its many hypophosphorylated peptides. The protein changes are reflected by similar transcript upregulation for SQSTM1, versus downregulation for SPARCL1, but the MYO6 protein accumulation is counteracted by downregulation of *Myo6* mRNA - a possible correlate of deficient autophagosome fusion with lysosomes. * *p* < 0.05; ** *p* < 0.01, *** *p* < 0.001, **** *p* < 0.0001, ns = not significant.

Overall, this study of spinal cord tissue in our SCA2/ALS13 mouse model documented massive inductions for SQSTM1 and NBR1 proteins, which are known to accumulate in TDP-43 aggregates upon ALS-associated mutations [118–122]. Conversely, our spinal cord study observed a loss-of-function of OPTN where autosomal recessive mutations are known to cause ALS type 12 [123–128]. Subsequent analyses of autophagy mechanisms *in vitro* focused on the regulation of these factors.

Interestingly, the induction of SQSTM1 protein was >5-fold. This suggests that several increases of phospho-peptides in SQSTM1 can be explained by higher protein abundance, only the massive increase at pSer152 near the ZZ domain, with the strong increases at residues 207-226 around the TRAF6-binding sequence (TBS), as well as residues 330-363 around LIR and Keap1-interacting region (KIR) reflect selective phosphorylation efforts. The *Sqstm1* mRNA was induced only <1.5-fold, suggesting that multiple translation rounds occur to amplify SQSTM1 availability. Asking whether ATXN2 as a factor known to influence mRNA stability and translation efficiency has a selective impact on this event, we interrogated previously published data [57] on the identity of mRNAs that are bound directly by ATXN2. As shown in Fig. S4, a specific interaction of Ataxin- 2 to *Sqstm1* transcript was reported in its 3’ untranslated region (3’UTR), overlapping with sequences binding to the RNA helicase DDX6, and with an interaction site for miR-183-5p. This microRNA is known to regulate the levels of SQSTM1 and to have abnormal levels in ALS [129, 130]. Thus, ATXN2 might influence the levels of SQSTM1 protein via a PRM-SH3 bridge, and the stability/translation of *Sqstm1* mRNA via its 3’UTR.

### 2.4. Regulation of autophagy factors upon BafA1 and NaARS administration, in embryonic fibroblasts from Atxn2-CAG100-KIN mice and skin fibroblasts from SCA2 patients

Fibroblasts from the mouse model and from SCA2 patients were used to test whether the regulation of autophagy factors in response to stressors is altered by ATXN2-polyQ expansions, using alternatively the drug Bafilomycin A1 (BafA1) that inhibits autophagy by interference with lysosome acidification [131], or the drug Sodium Arsenite (NaARS) that triggers protein misfolding via oxidative stress and thus promotes stress granule formation in parallel to protein breakdown [132].

The interference with autophagic degradation by BafA1 treatment showed the following effects in mouse embryonic fibroblasts (MEF): It impaired autophagic flux as detected by LC3-II accumulation at higher dosage both in wildtype and mutant cells, but the KIN cells showed consistently much less soluble ATXN2 protein levels, impaired inducibility of SQSTM1 protein, and tendentially lower OPTN abundance (Fig. 4A/B).

**Figure 4.**
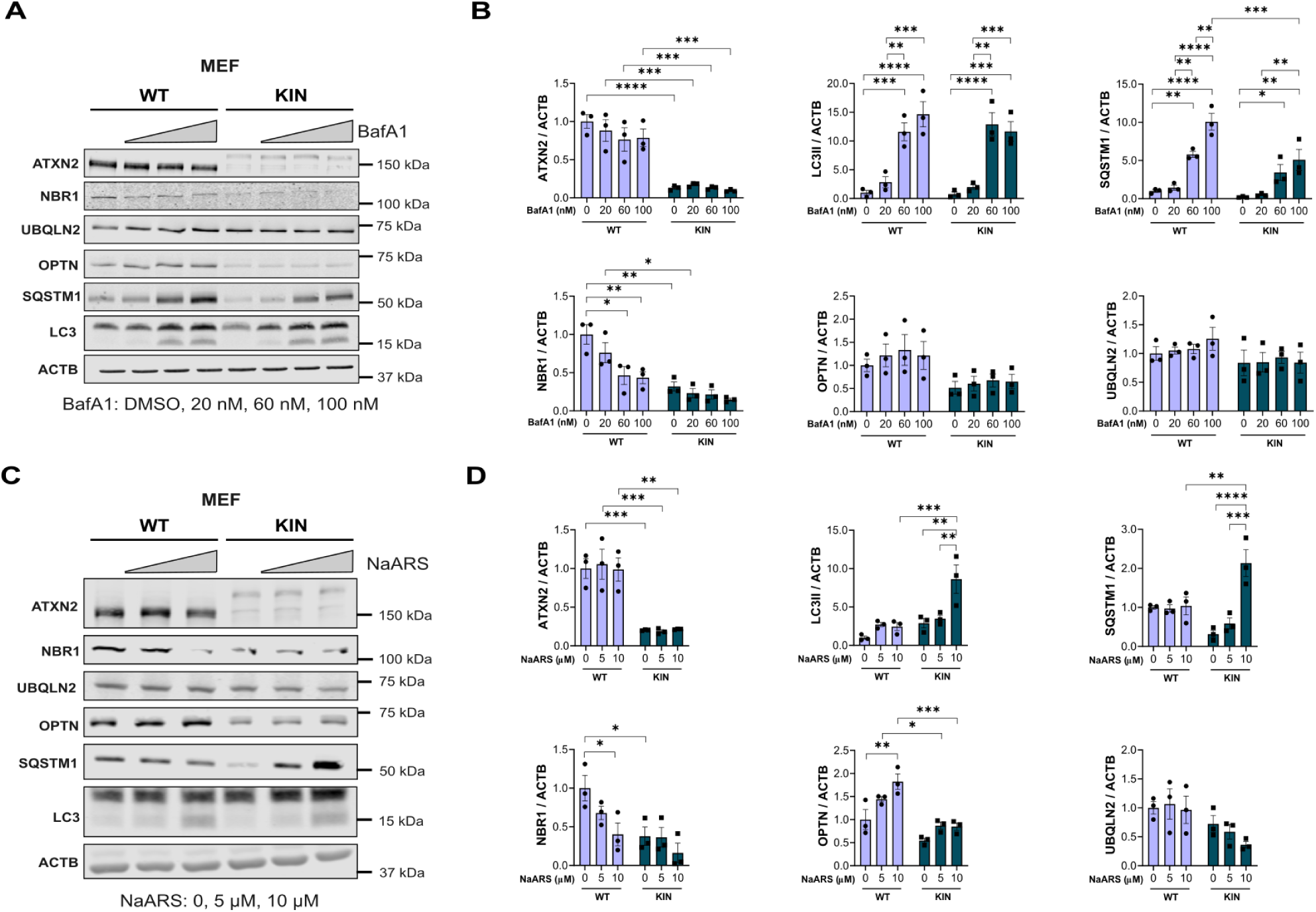
Quantitative immunoblots of autophagy factors in *Atxn2*-CAG100-KIN (KIN) versus wildtype (WT) MEF (N = 3 vs. 3) treated at increasing dosage either **(A)** with the the lysosomal toxin Bafilomycin A1 (BafA1) or **(B)** with the oxidative stressor Sodium Arsenite (NaARS). * p < 0.05; ** p < 0.01, *** p < 0.001, **** p < 0.0001.

The generation of high autophagy demand by NaARS treatment generated overlapping effects: In WT MEF, it did not affect LC3-II or SQSTM1 levels, while NBR1 decreased and OPTN increased with dosage. In KIN MEF, NaARS caused accumulation of LC3-II (>8-fold) and SQSTM1 (>2-fold), while the abundance of NBR1 and OPTN showed genotype-dependent reductions (Fig. 4C/D).

Overall, these data agree with the OPTN deficiency that was observed in KIN spinal cord, and suggest the SQSTM1 induction to be deficient in SCA2/ALS13. OPTN induction shows a smaller size than SQSTM1 induction, underscoring their differing importance for aggrephagy.

Next, three previously described SCA2 patient fibroblasts (polyQ expansions to CAG40/41/49, females with manifestation ages at 17-53 years [67]) versus sex-/age-matched controls were treated with BafA1. ATXN2 abundance exhibited almost normal levels in these human peripheral cells, presumably because the expansion sizes were much smaller than in the KIN MEF. The BafA1 treatment caused impaired autophagic flux as detected by more LC3-II abundance at higher drug dosage both in wildtype and mutant cells (less in SCA2). Selectively the SCA2 cells showed the accumulation of soluble LC3-II and SQSTM1 proteins to be restricted at high dosage, as previously observed for LC3-II in ATXN2-Q58 KnockIn HEK293 cells [133], in parallel to higher NBR1 abundance and trends to elevated UBQLN2 abundance (Fig. 5A/B). The findings may be explained by LC3-II and SQSTM1 becoming partially insoluble, and suggest that autophagy problems are triggers for the aggregation process. Furthermore, the data reveal cellular efforts in human SCA2 for a putative compensation via NBR1-dependent aggrephagy and UBQLN2-dependent ubiquitin/proteasome-mediated proteolysis.

**Figure 5.**
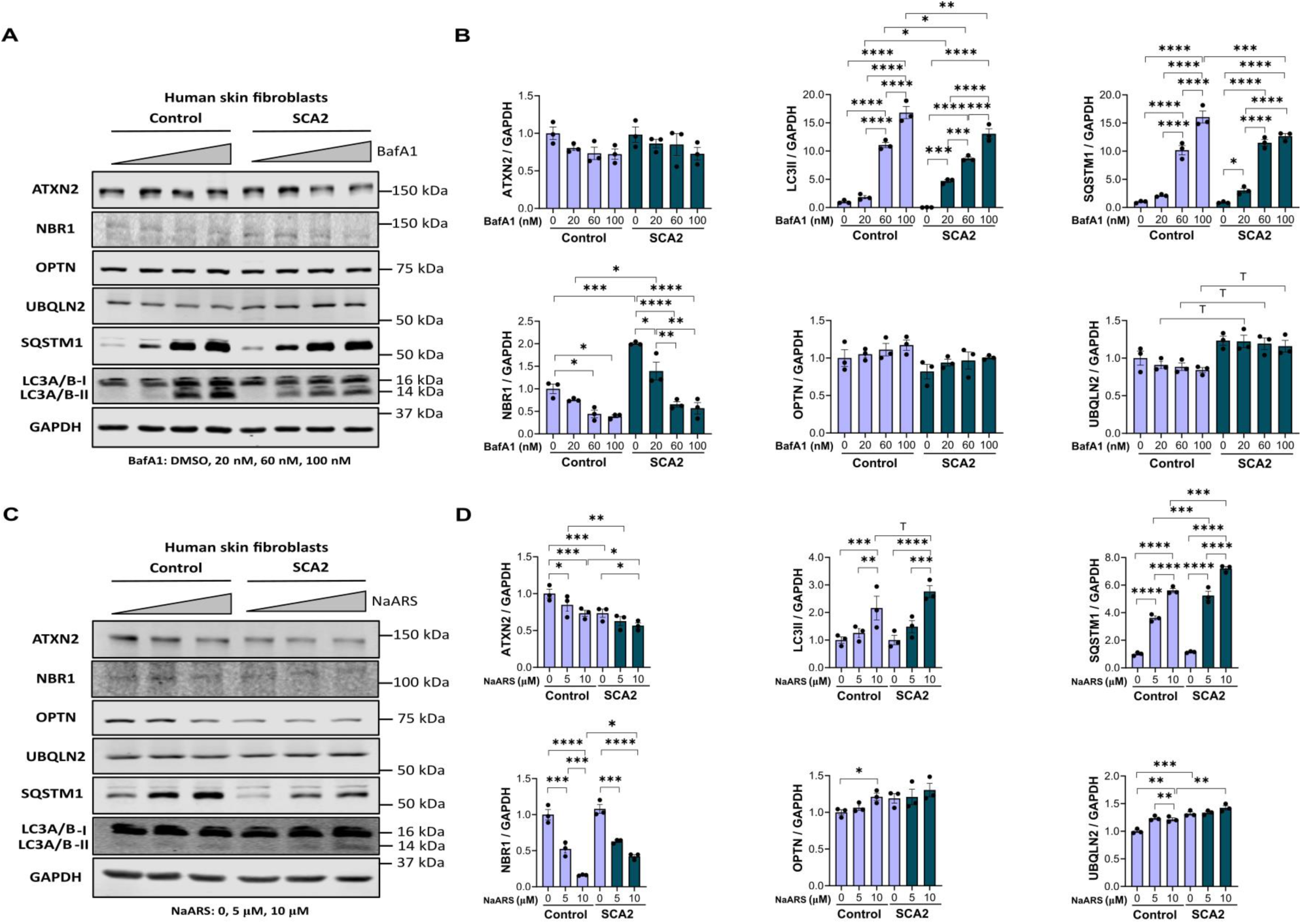
Quantitative immunoblots of autophagy factors in SCA2 patient skin fibroblasts (N = 3 mutants versus 3 controls) treated at increasing dosage either **(A)** with the lysosomal toxin Bafilomycin A1 (BafA1) or **(B)** with the oxidative stressor Sodium Arsenite (NaARS). T < 0.10; *p < 0.05; ** p < 0.01, *** p < 0.001, **** p < 0.0001.

Upon NaARS treatment of human skin fibroblasts with high demand for autophagic proteolysis, the accumulation of SQSTM1 and LC3-II was elevated in SCA2 patient cells, suggesting that the inducibility of SQSTM1 and LC3-II is not limited by the typical polyQ expansion sizes in most patients (Fig. 5C/D). Again, compensatory increases of NBR1 and UBQLN2 were found.

It was interesting to note that aggrephagy appeared to enhance the need for SQSTM1 while lowering endogenous NBR1 levels, suggesting that they can represent alternative SAR options, an observation that contrasts with the mutual positive regulation of both factors reported during viral infection and for adenovirally transfected NBR1 [134, 135]. Overall, both *in vitro* models of SCA2/ALS13 indicate that the polyQ expansion enhances the cellular need of SQSTM1 and autophagic flux upon exidative stress, but restricts the maximal availability of soluble SQSTM1 protein. These observations might be explained by LLPS of expanded ATXN2 with SQSTM1 into aggresomes, even in peripheral fibroblast cells.

### 2.5. Subcellular localization of polyQ-expanded ATXN2 and SQSTM1 positive aggresomes in Atxn2-CAG100-KIN spinal cord

To characterize the aggresome formation in SCA2 by an additional approach, triple immunofluorescent immunohistochemistry was employed to document the cytoplasmic localization of ATXN2 and SQSTM1 in affected neurons of the spinal cord (Fig. 6), in comparison to PABPC1 as a known direct ribonucleoprotein interactor of ATXN2 [44, 50, 136]. A previous publication already documented the sequestration of ATXN2 and TDP-43 to aggregates in spinal motor neurons of aged KIN mice [81]. In the current study, SQSTM1 aggregates were readily apparent in the motor neuron cytosol of KIN mice at opposite ends of the nucleus where aggresomes form near microtubule organizing centers (MTOC), as previously observed [137] (Fig. 6, upper row). While ATXN2 also showed aggregate formation at opposite ends of the nucleus in spinal motor neurons of KIN mice, PABPC1 staining showed mostly a diffuse cytoplasmic localization (Fig. 6, lower row). Thus, in KIN spinal cord motor neurons the polyQ-expanded ATXN2 and SQSTM1 show enhanced transport via the cytoskeleton to MTOCs.

**Figure 6.**
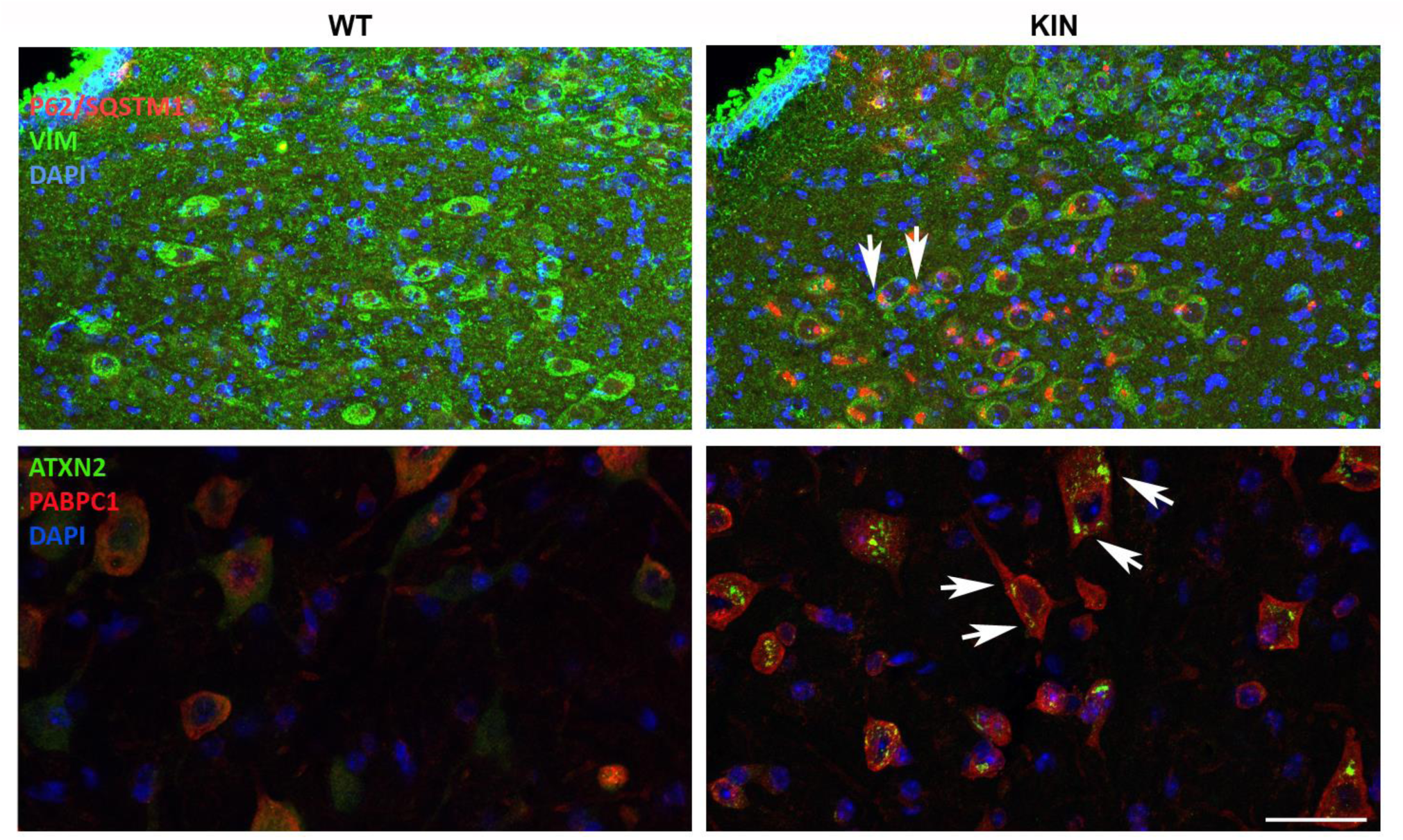
Triple immunofluorescent histochemistry by confocal laser scanning microscopy with composite panels from Z-stacks. Detection was done for the selective autophagy receptor SQSTM1 and the ribonucleoprotein PABPC1, compared to intermediate filament vimentin (VIM) or ATXN2, and nuclear DAPI signals, in the spinal cord of old sex-/age-matched adult wildtype (WT) versus *Atxn2*-CAG100-KnockIn (KIN) mice. White arrows in the upper row point to SQSTM1 aggregates at opposite poles of the nucleus, where microtubule organizing centers are expected to establish neuronal polarity and provide a starting point for axonal or dendritic transport. White arrows in the lower row point to ATXN2 aggregates at opposite poles of the nucleus.

### 2.5. Coimmunoprecipitation of ATXN2 from cerebellum with identification of interactors reveals binding increase for SH3-containing myosins, but reduction for ribonucleoproteins in KIN mice

To further understand the underlying protein-protein-interactions of soluble ATXN2, coimmunoprecipitations (CoIP) with subsequent mass spectrometry were performed. Given that spinal cord is composed to a large part of fiber tracts with myelinated axon bundles, where ATXN2 would be present only in minimal amounts, and in view of the unavailability of sufficient spinal cord tissue for maximizing sequencing depth, these analyses were performed in cerebellar tissues. As negative control of unspecific background signals, *Atxn2*-KO tissues were studied in parallel, although the main focus was on the comparison of KIN versus WT binding data. As expected, the comparison of WT to KO prominently identified the known ATXN2 interactions with ribonucleoproteins such as DDX6, LSM12, TARDBP (aka TDP-43) and others like PABPC1 and STAU1, with a clear ranking of their abundance ratios within the ATXN2 interactome (Table 2, Gene Symbols in blue letters).

**Table 2.**
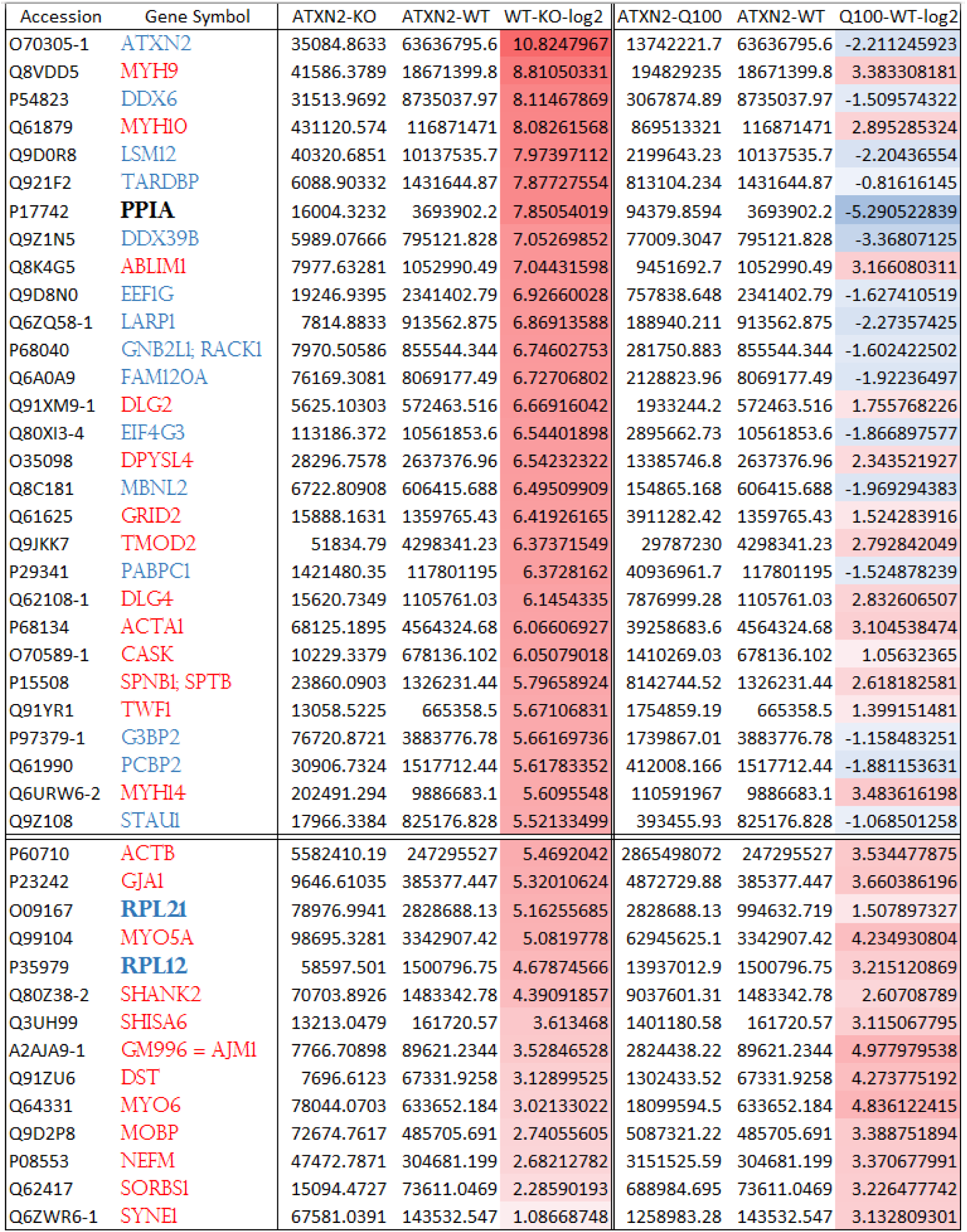
List of strongest ATXN2 interactors in mouse cerebellum. Accession column reflects UniProt database IDs. Ribonucleoproteins are shown as Gene Symbols with blue letters, while actomyosin-cytoskeleton-associated factors are in red letters. WT versus KO log2 abundance ratios (middle column) and KIN versus WT log2 abundance ratios (column at right margin) are illustrated as heatmaps with red color for increases, blue color for decreases. Usually ribonucleoproteins showed lower CoIP presence upon Q100 expansion, while cytoskeleton-associated factors showed enhanced presence, with significant enrichment of SH3-domain containing proteins among them.

A novel insight in Table 2 was the strong binding of ATXN2 to PPIA (peptidyl prolyl isomerase A). PPIA governs the function of TDP-43 for the assembly of ribonucleoprotein complexes [138], and the loss of PPIA in mice causes a neurodegenerative disease with key behavioral features of FTD, marked TDP-43 pathology and late-onset motor dysfunction [139]. A PPIA loss-of-function mutation was identified in a patient with sporadic ALS [139]. Drug inhibition of extracellular PPIA rescued motor neurons and extended survival in an ALS mouse model [140]. It was always unclear how cytosolic ATXN2 and nuclear TDP-43 are brought to interact, so the presence of PPIA in both subcellular compartments may contribute to this association.

Importantly, comparisons of KIN versus WT interactor abundance in the CoIP (Table 2, Table S2A/B) showed decreases for ribonucleoproteins (also PPIA), suggesting that reduced abundance and/or impaired affinity of polyQ-expanded ATXN2 with RNA-processing enzymes contribute to disease pathogenesis in SCA2/ALS via a partial loss-of-function mechanism.

As additional new interactors in the WT versus KO list, many actin-associated factors were prominent in the cerebellar CoIP (Table 2, Gene Symbols in red letters), in agreement with a previous report that mutations of Ataxin-2 in *Drosophila melanogaster* affect the actin cytoskeleton [75].

Myosin motor proteins and postsynaptic scaffolds were frequent among these interactors, with a prominent enrichment of (i) myosins with SH3-like domains in their N-terminus (MYO6, MYO5A, MYH9, MYH10, MYH14), (ii) postsynaptic SH3 domain proteins such as DLG4/2 or SHANK2, and (iii) the SH3-like domain containing ribosomal factor RPL21 (Fig. 7). In the KIN cerebellum, the abundance ratios increased consistently for these factors, in complete contrast to the decreases of ribonucleproteins. Crucially, these data validate the SH3 domain enrichment in the spinal cord phosphoproteome, and provide confirmatory evidence with a second approach that abnormal PRM-SH3-interactions contribute to the neurotoxic gain-of-function of Q100-ATXN2.

**Figure 7.**
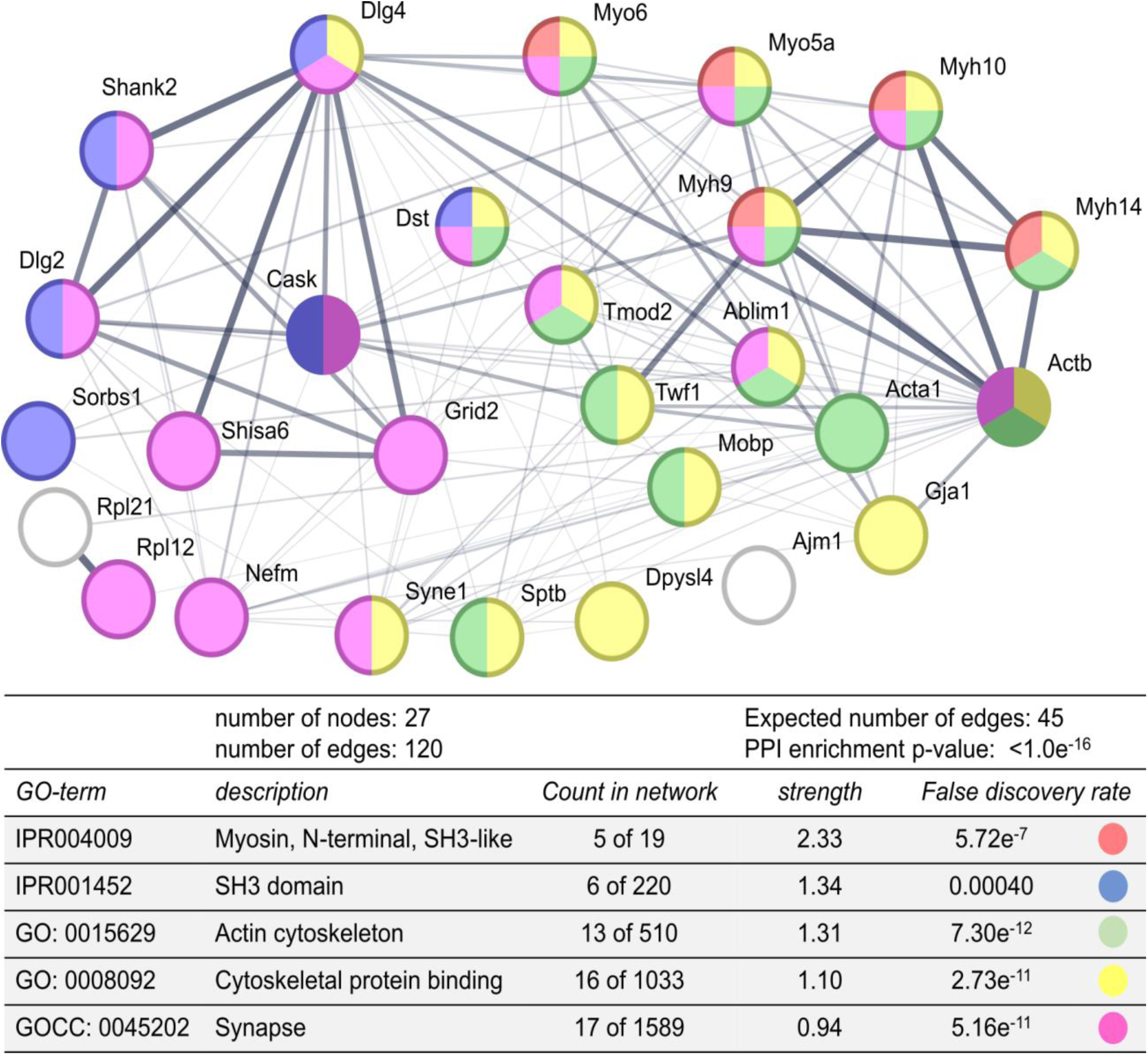
Significant enrichments of SH3 domain superfamily proteins, actin cytoskeleton, and synaptic machinery was observed upon STRING statistics among the ATXN2 interactors that showed a gain of binding upon polyQ expansion.

There were only two ribonucleoproteins with a similarly increased binding to polyQ expanded ATXN2, namely RPL12 and RPL21. This finding is in agreement with a publication where the yeast ortholog of ATXN2, named PBP1, was observed to regulate cell growth via association with RPL12 / uL11 [141]. RPL12 / uL11 forms the basis of the ribosomal P-stalk, where translation initiation, elongation, and fidelity are ensured, particularly during mitosis [142–144]. At the P-stalk, the EIF-2-Alpha Kinase GCN2 docks to couple translation with nutritional stress [145–149], and to regulate synaptic plasticity as well as neurite outgrowth [150, 151]. RPL21 is a member of the SH3-like domain superfamily, see UniProt entry O09167.

To enable a detailed analysis of these observations, Table S2 documents the cerebellar CoIP data, firstly for individual proteins with their label-free quantity per genotype (Table S2A/B), secondly for binding ratios per protein upon genotype comparison (Table S2C-E), thirdly for STRING enrichment statistics on proteins that specifically bind wildtype ATXN2 (Table S2F), fourth for increases or decreases in their association upon polyQ expansion (Table S2F-H), and fifth for enrichments of SH3-like domains in N-terminal myosin motors (FDR=5.72e-07), SH3 domain proteins (FDR=0.00040), actin-associated factors (FDR=2.73e-11), and synaptic machinery (FDR=7.14e-11) (Table S2I).

The SH3-like domains in the N-terminus of unconventional myosins provide actin-association. In particular, while the N-terminal SH3-domain in MYO6 links its motor domain to the cytoskeleton, the C-terminal MYO6 tail links it to TOM1 and autophagy receptors, such as OPTN and TAX1BP1 [152–156], ensuring the removal of aggregates. Thus, the gain of polyQ expanded ATXN2 binding to SH3-containing myosins like MYO6 might simply reflect its increased transport along the cytoskeleton to the autophagic degradation machinery. However, these observations are more likely explained by a MYO6 pathological accumulation due to a transport block, in view of the *Myo6* transcript downregulation (Fig. 3). Probably due to impossible solubilization of aggresomes for CoIP and mass spectrometry analysis, the enhanced binding of ATXN2 to SQSTM1 and NBR1 was not detected by this approach.

Overall, the CoIP survey confirmed a polyQ-expansion-triggered increase in the binding of SH3-containing proteins, especially for the actomyosin cytoskeleton that enables cell protrusion motility, and for the stimulus-response machinery in postsynapses.

## 3. Discussion

The molecular scenario of SCA2/ALS13 detailed above has strong similarity with the known polyQ expansion-driven neurodegenerative processes in autosomal dominant Huntington’s disease (HD). In HD, (i) exon-1 of the Huntingtin gene drives pathogenesis, (ii) erroneous splicing of this exon-1 and (iii) abnormal proteolysis of the N-terminal fragment were observed in KnockIn mouse models and patient cells, (iv) a proline-rich sequence lies adjacent to the polyQ domain, (v) its abnormal interaction with SH3 domains is responsible for the disease progression, by (vi) exerting selective aggrephagy effects via OPTN and MYO6 [152, 157–178].

In our study on SCA2/ALS13, the main finding is that the cellular phosphorylation responses to ATXN2 polyQ expansion appear massively and selectively upregulated within the N-terminus between residue Ser121 and Ser228. This observation suggests several important insights. (i) The possibility exists that an N-terminal fragment of polyQ-expanded ATXN2 is generated by abnormal splicing, translation, or proteolysis, to then accumulate in postmitotic cells, as driver of LLPS and the neurotoxic aggregation process. This fragment may be too insoluble and too small for unequivocal detection in immunoblots (the corresponding efforts were unsuccessful in our hands), but may have been present in the mass spectrometry [179]. Consequently, the antibodies against the polyQ-expansion domain (such as 1C2) and the antibodies against these N-terminal ATXN2 sequences would detect the aggregates much better than the current commercial antibodies raised against the remainder of ATXN2. Furthermore, it is crucial to consider how efficiently ASO are targeting and completely degrading the 5’-terminal sequences of the Ataxin-2 mRNA, for optimal prevention of the neurodegenerative process in SCA2/ALS patients. (ii) The neurotoxic gain-of-function may be determined by abnormal folding, binding, and solubility of the expanded polyQ domain and adjacent PRM domains of ATXN2 and its corresponding SH3-domain containing interactor proteins, affecting the actomyosin cytoskeleton together with the motility of cell membranes, in particular RTK and nutrient endocytosis, as well as ribosomal translation, resulting in the progressive atrophy of the nervous system. (iii) Crucially in the case of polyQ-expanded ATXN2, the phosphorylome and CoIP profiles suggest excessive autophagy in SCA2/ALS13 not to be an indirect late consequence in the course of disease, but a primary autodestructive phenomenon that is triggered by abnormal competition between ATXN2-PRMs and SQSTM1-PRM and/or WNK1-PRMs for association with SH3-domain proteins such as MYO6 as autophagosome motor protein, ITSN1/CIN85/SRC in endocytosis, RPL21 in ribosomal translation, FYN in myelination, and SHANK1 in glutamatergic postsynapses. As previously shown, the recruitment of SH3-containing proteins into phase-separated sequestosomes reduces their biological activity and function, e.g. in the case of SRC restricting the activation of PI3K and the formation of filopodia-like cell protrusions [180]. (iv) The phosphorylation changes reveal the primary switches where cellular compensation efforts occur. Their detailed analysis may guide researchers to the pathways where damage responses are impaired or insufficient. Such molecular events may be key targets for pharmacological efforts to prevent or postpone the clinical onset of motor deficits. (v) Apparently the physiological functions of ATXN2 and ATXN2L are adjusted with minor phosphorylation changes, and also other ribonucleoproteins do not appear altered among the prominent phospho-sites. The strong binding of WT ATXN2 to PPIA and their strongly decreased association upon polyQ expansion stands out as a novel finding that may explain the impact of ATXN2 on the ALS protein TDP-43. Overall, the phosphorylome and coimmunoprecipitation profiles of end-stage *Atxn2*-CAG100-KIN nervous tissues provided unprecedented insights where abnormal molecular interactions and signaling cascades exist that explain the characteristic disease features.

### 3.1. The N-terminus of ATXN2 with its polyQ repeat and three PRMs has a crucial role

As a result of alternative splicing [181], the main isoform of human ATXN2 begins with the sequence MSLKPQQQQQQQQQQQQQQQQQQQQQQQPPP (UniProt entry A0A2R8Y5A6), and the RNA-binding LSm domain starts 80 residues further downstream. The minor isoform includes additional 160 residues at the N-terminus (Uni-Prot Q99700), which contain three putative PRMs (firstly, amino acids 55-71 PPPPGPGPPPSRQSSPP, which appear to be responsible for most SH3 interactions according to Fig. 2 and Table S1L; secondly, aa 117-123 PPAAPTR; thirdly, aa 142-148 RPAPGCP, whose sequence would interact only with the SH3 protein NCF2 in competition with the PRM protein CYBA according to Table S1L). These sequences are highly conserved among mammals, in mouse only aa 117-123 has a relevant change to PAATPAR (UniProt O70305). This extended N-terminus does not exist in yeast, worms, flies, or zebrafish. During phylogenesis, the first N-terminal PRMs appeared in the ATXN2/ATXN2L ortholog of lizards (UniProt A0A670IJW1). Given that interactions of ATXN2 with SH3 domains will influence the actomyosin cytoskeleton below cell membranes, the ATXN2 isoform with extended N-terminus may have developed to ensure RNA quality during long-distance trafficking along the dendrites of larger animals, and particularly in peripheral cell compartments like synapses. In the cell periphery, ATXN2 with extended N-terminus and multiple SH3 interactions will also be more efficient in regulating the RTK-dependent endocytic uptake of nutrients, mTORC1 signaling, and stress/stimulus-dependent local translation. Thus, it will be impossible to faithfully model the diverse aspects of SCA2/ALS13 pathogenesis e.g. in yeast and flies, where the isoform with PRMs in the extended N-terminus does not exist.

### 3.2. Characteristic features of SCA2/ALS13 can be explained by preferential affection of SH3-PRM-containing proteins with conspicuous phosphorylation changes

Indeed, polyQ-expanded ATXN2 triggers phosphorylation changes with significant enrichment among SH3 proteins, which modulate actin and endocytosis dynamics in the cell periphery as well as ribosomal translation, and which are credible correlates of the SCA2/ALS-typical neural atrophy pattern. The preferential affection of glutamatergic synapses between upper and lower motor neurons of the corticospinal projection, and between granule neuron axons known as parallel fibers and Purkinje neuron dendrites, could be explained by polyQ-ATXN2-triggered phase separation of SH3-containing proteins, e.g. the hyperphosphorylated SHANK1 and the enhanced interactors SHANK2 and DLG4 (aka PSD-95) as postsynaptic components of excitatory synapses [182]. LLPS of SHANK might underlie the tandem hypophosphorylations in the PRM-containing metabotropic glutamate receptor GRM1 (aka MGLUR1) as main factor for dendritic spine biogenesis and as disease protein responsible for autosomal recessive SCAR13 and autosomal dominant SCA44 [183, 184], and might underlie also the prominent multiple hypophosphorylations / reduced abundance of SPARCL1 as a factor responsible for the formation of excitatory synapses [113]. An association with the pathogenesis of motor neuron disease was previously reported for SPARCL1 (aka Hevin) [117, 185, 186]. Also the strong demyelination in SCA2 can be correlated to the hyperphosphorylation of the SH3-containing protein FYN as main regulator of axon-oligodendroglia signaling [187] with its interactor protein FYB1 / ADAP [188], and to adaptive, prominent, multiple hypophosphorylations in the PRM-containing myelin proteins MAG, MBP and MOBP [189]. As seen in Fig. 2, many other interactions between SH3 and PRM proteins show mild hyperphosphorylations, suggesting they might be partially affected by the phase separation process, e.g. SH3 proteins DLG1, CTTN and TJP1 / ZO-1 as well as PRM proteins AFDN, PXN, VASP and PAK3 as modulators of synaptic adhesion [190–196], or SH3 protein NCF2 and PRM protein CYBA as mediators of microglial neuroinflammation [197].

A further combination of hyperphosphorylations in the SH3 proteins ITSN1, SRC, SH3KBP1 / CIN85 as endocytosis regulators, versus hypophosphorylations in the PRM protein ARHGAP32 / PX-RIX as endosome-associated factor [198] was observed for the nutrient uptake pathway that governs mTORC1 signaling and cell growth. Finally, the combination of hyperphosphorylation in the SH3 protein AHI1 as endocytosis/autophagy modulator [199], as well as accumulation of the SH3 protein MYO6 as autophagosome maturation factor [152], versus multiple prominent hypophosphorylations / reduced abundance in the PRM-containing proteins (Table S1K) WNK1 as autophagy regulator [200], ATG2A as autophagy initiation factor [201], and TOM1L2 as xenophagy mediator [202] also suggests these events to be among the primary erroneous protein interaction consequences upon polyQ expansion in ATXN2. The enhanced binding of Q100-ATXN2 to the ribosomal components RPL12 and the SH3-containing RPL21 may impair initiation, elongation and fidelity of mRNA translation, thus contributing to the mTORC1 inhibition phenotype.

In view of the importance of PRM-SH3-interactions for the neurotoxicity of polyQ expansions in SCA2/ALS13, it should be noted that targeting the SH3-containing SRC protein and its signaling pathway was recently proposed to develop a preventive treatment of ALS [203].

### 3.3. Excess autophagy may be a primary direct effect, leading to SQSTM1 and OPTN deficiciency

A direct effect of polyQ-expanded ATXN2 on autophagy can be caused by four parallel mechanisms, which are compatible with the data observed. Firstly, LLPS of PRM-containing WNK1 and its multiple hypophosphorylations may trigger its activity loss. Secondly, abnormal interactions of the PRM and SH3 domain in hyperphosphorylated TP53BP2 / ASPP2 may modify its autophagy initiation role [204, 205]. Thirdly and perhaps as a critical ALS-associated mechanism, a partial deficiency of OPTN may be caused by its association with PRM-containing hyperphosphorylated TOM1 and hypophosphorylated TOM1L2 (Table S1K), and its association with accumulated SH3-containing MYO6. Fourth and perhaps most importantly among ALS-associated mechanisms, the massive hyperphosphorylation of PRM-containing SQSTM1 may reflect the outstanding cellular efforts to promote this aggrephagy process. This compensatory mechanism may parallel its exceptionally rapid inducibility after oxidative stress by NaARS, but contrasts with its restricted availability upon autophagy interference by BafA1 in SCA2 cell models. Given that ATXN2 was observed to bind *Sqstm1* mRNA directly, and that SQSTM1 acts as the selective autophagy receptor that mediates the trafficking of phase-separated polyQ-expanded ATXN2 to MTOC-associated aggregates and to subsequent proteolytic degradation via autophagosomes/lysosomes, this pathway may contain key targets for neuroprotective therapy approaches.

### 3.4. Phospho-site regulation of protein activity in autophagy and proteostasis

The strong hyperphosphorylation events in proteostasis factors may provide clues about regulatory mechanisms, which may become useful in the development of neuroprotective therapies. Certainly, it is important to consider in detail whether phosphorylation events such as pSer152-SQSTM1, pSer207/Ser224/Ser226-SQSTM1, pSer344/Ser351/Ser357/Ser363-SQSTM1, pSer547-OPTN represent an ALS-specific phospho-profile among the SAR of aggrephagy, and pSer267/Ser271-UBQLN2 an ALS specific phospho-profile of this misfolded protein delivery factor to proteasome or/and autophagy.

SQSTM1 with its 5-fold increased abundance is the primary interest for researchers into the aggrephagy mechanisms in SCA2 and ALS, given that its mutation can result in a neurodegenerative process with the phenotype of ALS type 3 [122]. Its 107.6-fold hyperphosphorylation within the ZZ domain (as cargo binding sequence) at pSer152-SQSTM1 was previously attributed to TBK1 [206], a Ser/Thr kinase whose mutation causes ALS [207]. In addition, three clusters of phospho-sites were located within an intrinsically disordered region. Firstly, three phospho-sites between amino acids 207 and 226 (pSer207 under control of RNF166 as autophagy regulator [208]) with on average 36.7-fold increases were clustered in the vicinity of the TBS motif, where TRAF6 binding modulates the neurotrophin signaling cascades [209]. Secondly, three phospho-sites between amino acids 334 and 344 with on average 42.7-fold increases were clustered in the vicinity of the LIR motif, where LC3 binding determines the association of SQSTM1 with autophagosome membranes [209]. Thirdly, four phospho-sites between amino acids 351 and 363 with on average 32.9-fold increases were clustered in the vicinity of the KIR motif, where KEAP1 binding modulates the NRF2-dependent transcription of detoxifying, cytoprotective, antioxidant and inflammation-modulating factors [209]. Thus, the SQSTM1 hyperphosphorylations may alter not only the association with autophagosomes, but also with aggregate cargo and with cellular types of machinery for trophic signaling and stress responses. They are located in areas that explain the altered stress responses preferentially for nervous tissue, and may constitute an ALS-specific pattern within pathogenesis.

Despite its decreased abundance, OPTN showed a 2.7-fold increase of pSer187 within the LIR domain, which regulates the binding to autophagosome membranes, and a 10.0-fold increase of pSer547 close to the ZF motif and within the binding area for the ER-to-Golgi vesicle traffic regulator RAB1A, and for TAX1 in competition with TAX1BP1 [113, 114]. As mentioned, OPTN mutations cause ALS type 12 [210]. Furthermore, OPTN is regulated by the ALS-associated kinase TBK1 [211, 212].

The 10.7-fold increase of pSer313-TAX1BP1 is in the vicinity of the O homodimerization motif (amino acid 320-340) [213], and between the coiled-coil CC1 domain (amino acid 154-307) and the CC2 domain (amino acid 349-476) that is responsible for self-oligomerization [214]. So far, TAX1BP1 was not found to be associated with selective neurodegeneration phenotypes such as ALS or ataxia, but it is targeted by the ALS-associated kinase TBK1 and interacts with the TBK1-regulated putative ALS disease protein and neuroinflammation repressor TNIP1 (aka ABIN1). TNIP1 inhibits selective autophagy via competition with SARs, and TNIP1 abundance is controlled by autophagy [215–222]. A 2.7-fold hyperphosphorylation at pSer279-TNIP1 implicates this factor in SCA2/ALS13 pathogenesis. Apparently, TAX1BP1 plays its main role in the amplification of the aggregation process and the maturation of autophagosomes.

NBR1 as the archetypal SAR in aggrephagy [223, 224] is not associated with ALS. It showed >2.5-fold increased abundance in KIN spinal cord, appeared regulated in reciprocal relation to SQSTM1 in cultured cells, and its KIN spinal cord phospho-profile showed increases in two areas. These hyperphosphorylations were 6.4-fold at pSer116 and 9.8-fold at pSer117 in proximity to the homo-multimerization domain PB1, and 2.2-fold at pThr647 between the LIR1 and LIR2 domains responsible for autophagosome membrane binding [224]. Thus, cargo-protein interactions were not modulated in contrast to SQSTM1 and OPTN.

Despite the physiological localization of ATXN2 with the rough endoplasmic reticulum (rER) [43] and its relocalization to stress granules after cell damage [58], no putative ribophagy or ER-phagy receptor (such as NUFIP1, FAM134B, RTN3L, SEC62, CCPG1, ATL3, TEX264, NIP-SNAP1/2) appeared hyperphosphorylated. Only the hyperphosphorylation of UBQLN2 with its ability to access phase-separated RNP complexes and to degrade rER proteins via the proteasome or via lysosomes [225], with an ALS phenotype upon mutation [226, 227], appeared to relate to the RNA binding features of ATXN2 and to its preferential impact on motor neurons. UBQLN2 hyperphosphorylations at pSer271 (17.2-fold) and pSer267 (14.6-fold) were located in the middle of the four STI1 domains, where the association with molecular chaperones is regulated [228]. Subsequent to chaperone interaction, UBQLN2 can either activate the ubiquitin-proteasome pathway of proteolysis, or promote autophago-lysosome formation. Both hyperphosphorylations are closest to the second STI1 domain, where LLPS is modulated and the relocalization to stress granules and ribonucleoprotein complexes is determined [229, 230], so that UBQLN2 can promote the disposal of rER proteins [225]. This preferential role for RNA associated protein elimination may underlie the specific strong link between UBQLN2 mutations and the ALS phenotype [228, 229, 231].

The hyperphosphorylation of HSPB1 may also reflect cellular efforts to promote refolding and chaperone-mediated autophagy, particularly of the actin cytoskeleton and RNA granules [232–234]. WDFY3 abundance was increased almost 3-fold in KIN spinal cord, so the 3.2-fold elevated presence of pThr818 may only reflect the accumulation of this activator of selective aggrephagy that cooperates with SQSTM1, LC3-II and ATG proteins [235–237]. This phospho-site lies within an Armadillo-like multi-helical fold (residues 460-940) that is crucial in the yeast autophagy factor Vac8 for the association with disordered extended C-termini of ATG proteins, the recruitment of cargo, and the link with membranes of the proteolytic vacuole [238].

The 8.7-fold increase of pSer216-CTSB is within the heavy chain area where several posttranslational regulations occur, at amino acid 211 as part of a disulfide bond, at amino acid 220 where an N6-acetyl-lysine modification was observed [239], and at amino acid 240 where S-nitrosylation controls enzyme activity [240]. CTSB can be secreted [241], but in lysosomes it also serves as apical regulator that is in control of overall lysosomal dynamics and autophagy [242]. Again, CTSB has no association with ALS, it seems to be an unspecific factor in autophago-lysosomal pathology.

In contrast, BNIP3 acts in mitophagy, exhibited a widespread non-selective phosphorylation increase of mild degree, and is not associated with ALS. BNIP3 abundance was increased >3-fold, so the elevated amounts of phospho-peptides (2.2-fold at pSer48, 2.3-fold at pSer79, and 2.4-fold at pSer85, pThr86, pSer88) may only be a secondary phenomenon. Altered mitophagy via BNIP3 adaptation to polyQ expansion may serve to explain the induction of *Pink1* mRNA as kinase responsible for mitochondrial degradation in the blood of SCA2 patients [243].

Overall, the cellular efforts to optimize autophagy for the degradation of polyQ-expanded ATXN2 aggregates involve phosphorylations and adjusted abundance of several proteostasis factors, namely (i) in the cargo recognition domain of SQSTM1, (ii) in the LIR motifs of SQSTM1 and OPTN, (iii) in the homo-multimerization domains of TAX1BP1 and NBR1, (iv) regarding chaperone-proteasome roles via UBQLN2 and HSP1B, (v) regarding lysosome availability via WDFY3 and CTSB. Elevated mitophagy may be a side-effect of polyQ-ATXN2 aggregate degradation.

### 3.5. Mild phosphorylation reductions in ATXN2 C-term, ATXN2L, ribonucleoproteins, and PPIA

The reduced appearance of phospho-sites in ATXN2 from pSer319 to the C-terminus upon polyQ expansion is probably a consequence of its impaired transcription and translation. Most of these hypophosphorylated sites are clustered within the IDR2 of ATXN2 (Fig. 1B), where microtubule binding is determined, and where a mild influence on mRNA and RNP binding was documented [49, 52, 244]. In view of the combination of a C-terminal partial loss-of-function versus an N-terminal neurotoxic gain-of-function, it was interesting to examine the regulation of its paralog ATXN2L. Previous studies in ATXN2L-mutant cells had not observed an adaptation of ATXN2 levels, and vice versa [37], so the functional relationship between the two gene copies had remained unclear. The ribonucleoprotein ATXN2L has higher levels than its paralog ATXN2 and exerts an essential role in embryonic development [37]. As a novel insight, a mild hyperphosphorylation (4.1-fold) of presumably compensatory nature occurred in ATXN2L at pSer333, within the IDR2 very close to the LSm-AD region (amino acids 262-331) (Fig. S1), where binding to RNA helicases like DDX6 is thought to impact the processing of mRNAs and the silencing of transcripts [42, 58–60]. This ATXN2L pSer333 change is known to be controlled by Protein Kinase D1 (PRKD1), as documented in the internet database at www.phosphositeplus.org (last accessed on 28 February 2024). This finding provides insight into the signaling mechanisms that govern ATXN2L physiology. It was previously established that ATXN2 via its proline-rich domains modulates the uptake of trophic factor receptors such as EGFR, via binding to SH3 motifs in endosome trafficking proteins such as SRC and GRB2 [65, 67], and that ATXN2 can localize to the trans-Golgi network [245]. Therefore, it is very noteworthy that PRKD1 also acts in the Golgi apparatus, converting transient diacylglycerol (DAG) signals to control Golgi integrity, establishing trafficking polarity, and delivering membrane proteins to neuronal dendrites for the maintenance of arborizations, in particular in cerebellar Purkinje cells [246–250]. PRKD1 also stabilizes actin in dendritic spines with consequences for synaptic plasticity and long-term memory [251], modulates the endocytosis of AMPA receptors in glutamatergic postsynapses [252], and confers neuroprotection against excitotoxicity [253–256]. Mutant PRKD1 leads to an age-dependent decline of locomotion rate in *Caenorhabditis elegans* nematode worms [257]. Beyond its well-documented roles in neurons, PRKD1 also acts in glial cells to control nutrient uptake via micropinocytosis, as well as protein aggregate uptake via phagocytosis [258]. Thus, the impact of deficient PRKD1 function has a close resemblance to the features of SCA2, and acts on similar subcellular compartments as ATXN2/ATXN2L.

With regard to other ribonucleproteins, it was conspicuous that they showed decreased presence in the ATXN2-CoIP upon polyQ expansion, and that cellular phospho-signaling to adjust their function was a minor feature in KIN tissue. Curiously, this was also true for TDP-43 as ALS disease protein and known ATXN2 interactor. It is noteworthy that our mass spectrometry data in Q100-KIN mouse spinal cord did not replicate the increase of pTDP-43 upon ATXN2 intermediate polyQ expansions, which was reported based on quantitative immunoblots in postmortem spinal cords from ALS cases [259]. Contrasting with ribonucleoproteins, their associated folding modulator PPIA showed decreased presence in ATXN2-CoIP from KIN cerebellum, but accumulation in immunoblots from KIN spinal cord possibly due to altered TDP-43 association, and accompanied by a relevant 4.1-fold increase of pSer159-PPIA. These observations suggest that cellular compensatory efforts against ALS pathogenesis are targeting this molecular switch. The location of this phospho-site in the PPIA C-terminal region is close to several N6-acetyllysine modifications at amino acid 125, where interaction with TDP-43 is modulated [138].

### 3.6. Are pathogenic or compensatory phosphorylation changes suitable targets for intervention?

Although the particularly massive hyperphosphorylations are at the center of our interest, and may constitute targets of novel neuroprotective approaches, a cautionary comment has to be made. Similarly striking findings were previously observed in protein aggregation-driven neurodegenerative disorders, as in the disease protein Tau (e.g. at its 233Pro-Lys-pSer-Pro motif) for Alzheimer’s disease and FTD [260–262], and in the disease protein TDP-43 (at its C-terminal 404pSer-pSer-Met-Asp-Ser-Lys-pSer-pSer motif) for ALS and FTD [263, 264]. However, it has proven very difficult to distinguish between protective versus toxic phosphorylation events; even for the hyperphosphorylation of TDP-43, the subsequent experiments have claimed either a compensatory role to prevent pathogenic TDP-43 isoforms from aggregation [264–267], or a contrary pathogenic role [268, 269], or a role according to residue and context [270, 271]. Clearly, hyperphosphorylation can provide a crucial clue to define, which proteins and polypeptides are most affected by aggregation, and where liquid-liquid phase separation (LLPS) would lead to massive conformational changes. However, the skillful adaptation of site-specific phosphorylation dosage to progressive stages of disease is currently difficult.

### 3.7. Conclusion

Novel insights regarding the ATXN2 interactome and the adjustment of cell phospho-signals upon Q100-ATXN2 expansion indicate a prominent role of abnormal PRM-SH3 interactions that affect endocytosis/autophagy, actomyosin cytoskeleton, and synaptic pathways directly. These pathways are probably crucial for efficacious preventive therapy.

## 4. Materials and Methods

### Experimental animals

All mouse experiments were in conformity with the German Animal Welfare Act, the Council Directive of 24th November 1986 (86/609/EWG) with Annex II, and the ETS123 (European Convention for the Protection of Vertebrate Animals). All mice were housed at the Central Animal Facility (ZFE) of Goethe University Medical School, kept in individually ventilated cages with nesting material at a 12 h light/12 h dark cycle, with appropriate temperature and humidity conditions, and provided with food and water *ad libitum*. Breeding of heterozygous carriers was used for colony expansion and maintenance. Among offspring littermates, the homozygous *Atxn2*-CAG100-KIN and WT animals of the same sex were selected and aged together in the same cages for subsequent experimental group comparisons regarding phosphoproteome, gene expression, protein abundance, and immunohistochemistry. Both female and male mice were used during all experiments. The study was ethically assessed by the Regierungspräsidium Darmstadt, with approval number V54-19c20/15-FK/1083.

### Genotyping of Atxn2-CAG100-KIN mice

The genotyping of *Atxn2*-CAG100-KIN mice was done as previously described [77]. Briefly, genomic DNA was isolated from ear punches or tail biopsies using a standard protocol [272]. The CAG repeat was assessed by standard polymerase chain reaction (PCR) amplification with the NOW1-K2 and NOW1-H2 oligonucleotide primers, TaKaRa LA Taq-Polymerase (Takara Bio Inc., Japan), and a PCR Master Mix containing 2 mM Cresol, 250 µM dNTPs, and 60% sucrose in PE buffer. Amplification was performed in a SimpliAmp^TM^ PCR thermocycler (Applied Biosystems, Foster City, CA, USA) as follows: initial denaturation at 94 °C for 3 min, followed by 30 cycles of denaturation at 94 °C for 15 min, annealing at 68 °C for 4 min, elongation at 68 °C for 4 min, and a final elongation step at 68 °C for 9 min. PCR products were run in 2% agarose gel electrophoresis and the TrackIt™ 100 bp DNA Ladder (Invitrogen, USA) was used to estimate the size of the amplicons. The WT allele produced an amplification product of 793 bp and the expanded CAG100 allele a 948 bp amplicon.

### Cell culture and treatments

MEF were obtained from WT and homozygous *Atxn2*-CAG100-KIN embryos around E15–18 as previously described [48]. Human skin fibroblasts were acquired by biopsing the forearm of three clinically and molecularly diagnosed SCA2 patients and three age- and gender-matched control individuals, see [67]. MEF and human skin fibroblasts were cultured in Dulbeccós modified Eagle medium (DMEM) (1X) with high glucose (4.5 g/L D-Glucose) and sodium pyruvate (0.11 g/L) (Thermo Fisher Scientific #21969), supplemented with 1% glutamine (Gibco #25030081), 10% FBS (Gibco #A3160802), and 1% penicillin/streptomycin (Gibco #15140-122) at 37 °C, 5% CO2, and 95% relative humidity. The fibroblasts were plated in 100 mm dishes and allowed to grow for at least two days until they reached 70-90% confluency, before being treated with Bafilomycin A1 (BafA1) (Sigma #19-148), sodium arsenite (NaARS) (Sigma #S7400) or their appropriate vehicles (0.5% DMSO or H_2_O, respectively). Fibroblasts were treated overnight (24 h) with BafA1 (20, 60 or 100 nM) dissolved in dimethyl sulfoxide (DMSO), or NaARS (5 or 10 μM) dissolved in media. Equal number of dishes were treated with the appropriate vehicle only. Cell pellets were obtained by trypsination and centrifugation.

### IMAC Phosphoproteomics

Mouse cervicothoracic spinal cord tissue was obtained after cervical dislocation, then immediately frozen and transported in liquid nitrogen. Phosphoproteome profiling of spinal cord tissue from four WT versus four KIN mice was performed at Cell Signaling Technology INC. using their PTM-Scan and Fe-IMAC services. Lysed and trypsin-digested samples were desalted over C18 columns. A combination of anti-phospho motif antibodies and Immobilized metal affinity chromatography (IMAC) was used for phospho-peptide enrichment. Peptides were loaded directly onto a 50 cm × 100 µm PicoFrit capillary column packed with C18 reversed-phase resin. The column was developed with a 150-minute linear acetonitrile gradient in 0.125% formic acid delivered at 280 nL/min. Immunoprecipitated peptides were run in LC-MS/MS on an Orbitrap™ Fusion Lumos Tribrid mass spectrometer using a top 20 DDA method. MS Parameter Settings: MS Run Time 168 min, MS1 Scan Range (300.0 – 1500.00), Top 20 MS/MS (Min Signal 500, Isolation Width 2.0, Normalized Coll. Energy 35.0, Ac-tivation-Q 0.250, Activation Time 20.0, Lock Mass 371.101237, Charge State Rejection Enabled, Charge State 1+ Rejected, Dynamic Exclusion Enabled, Repeat Count 1, Repeat Duration 35.0, Exclusion List Size 500, Exclusion Duration 40.0, Exclusion Mass Width Relative to Mass, Exclusion Mass Width 10 ppm). MS/MS spectra were evaluated using Comet [273] and the Core platform from Harvard University [274, 275]. Searches were performed against the most recent update of the Uniprot *Mus musculus* database with a mass accuracy of +/-20 ppm for precursor ions and 0.02 Da for-product ions. Results were filtered with mass accuracy of +/– 5 ppm on precursor ions and the presence of the intended motif.

### Quantitative reverse transcription polymerase chain reaction in spinal cord and fibroblasts

Total RNA isolation from spinal cord tissue and fibroblasts pellets was performed with TRIzol Reagent (Sigma Aldrich, USA) according to manufacturer’s instructions. Total RNA yield and purity were quantified using a Tecan Spark plate reader (Tecan Group Ltd, Switzerland) at 230, 260, and 280 nm, in a NanoQuant plate. cDNA synthesis was performed from 1 μg of total RNA template using the SuperScript IV VILO kit (Invitrogen, USA) according to the manufacturer’s instructions. Gene expression profiles were assessed by quantitative reverse transcription polymerase chain reaction (RT-qPCR) using a StepOnePlus^TM^ (96 well) Real-Time PCR System (Applied Biosystems, USA). RT-qPCRs were run in technical duplicates on cDNA from 25 ng total RNA, with 1 μl TaqMan® Assay, 10 μl FastStart Universal Probe Master 2× (Rox) Mix (Roche, Switzerland) and ddH_2_O up to 20 μl of total volume. The PCR cycling conditions were 50 °C for 2 min, 95 °C for 10 min, followed by 40 cycles of 95 °C for 15 min and 60 °C for 1 min. The gene expression TaqMan® assays (Thermo Fisher Scientific, Waltham, Massachusetts, USA) used for this study were: Bnip3 (Mm00833810_g1), Ctsb (Mm01310506_m1), Myo6 (Mm00500651_m1), Nbr1 (Mm01249798_m1), Optn (Mm01333245_m1), Pcbp1 (Mm00478712_s1), Ppia (Mm02342430_g1), Sparcl1 (Mm00447784_m1), Sqstm1 (Mm00448091_m1), Tax1bp1 (Mm00518038_m1),Ubqln2 (Mm00834570_s1), Wdfy3 (Mm00556380_m1)., and *Wnk1* (Mm01184014_m1, Mm01184007_m1, Mm01184011_m1), and *Wdfy3* (Mm00556380_m1). The data were analyzed via the 2^−ΔΔCt^ method [276], using *Tbp* (Mm00446973_m1) as housekeeping gene.

### Protein extraction and immunoblots

Mouse cervicothoracic spinal cord tissues were homogenized with a motor pestle in 5-10× weight/volume amount of RIPA buffer consisting of 50 mM Tris-HCl (pH 8.0), 150 mM NaCl, 2 mM EDTA, 1% Igepal CA-630 (Sigma Aldrich, USA), 0.5% sodium deoxycholate, 0.1% SDS, cOmplete™ Protease Inhibitor Cocktail (Roche, Switzerland), and Halt™ Phosphatase Inhibitor Cocktail (Thermo Fisher Scientific, Inc., USA). Similarly, MEF and human skin fibroblasts pellets were homogenized in RIPA buffer by pipetting. The resulting protein suspensions were sonicated, and protein concentration was determined in a Tecan Spark plate reader (Tecan Group Ltd, Switzerland) using a Pierce™ BCA protein assay kit (Thermo Fisher Scientific, Inc., USA). 15 to 25 μg of total proteins were mixed with 2× loading buffer consisting of 250 mM Tris-HCl pH7.4, 20% glycerol, 4% SDS, 10% 2-mercaptoethanol, and 0.005% bromophenol blue, incubated at 90 °C for 5 min, separated on 8-15% polyacrylamide gels at 120 Volts, and transferred to nitrocellulose membranes (0.2 µm) (Bio-Rad Laboratories, Inc., USA). The nitrocellulose membranes were blocked in 5% BSA/TBS-T, and incubated overnight at 4 °C with primary antibodies. Afterwards, the nitrocellulose membranes were incubated for 1 h at room temperature, with fluorescently labeled secondary IRDye® 800CW goat anti-mouse (LI-COR 926-32210, 1:10,000), IRDye® 800CW goat anti-rabbit (LI-COR 926-32211, 1:10,000), IRDye® 680RD goat anti-mouse (LI-COR 926-68070, 1:10,000) or IRDye® 680RD goat anti-rabbit (LI-COR 926-68071, 1:10,000). Membranes were scanned using an Odyssey® Classic Imager. Image visualization and quantification of signal intensities was performed using Image Studio^TM^ software (version 5.2) (LI-COR Biosciences, Ltd., UK). The following primary antibodies were used: ATXN2 (Proteintech 21776–1-AP, 1:1000), BNIP3 (Proteintech 68091-1-Ig, 1:1000), CTSB (Abcam ab214428, 1:1000), LC3 A/B (Cell Sig-naling 4108, 1:1000), NBR1 (Cell Signaling 9891, 1:500), OPTN (Proteintech 10837-1-AP, 1:1000), PPIA (Proteintech 10720-1-AP, 1:1000), SQSTM1/p62 (Cell Signaling 39749, 1:1000), TAX1BP1 (Proteintech 14424-1-AP, 1:1000), UBQLN2 (Novus Biologicals NBP2-25164, 1:1000), WDFY3 (Proteintech 55009-1-AP, 1:1000), WNK1 (Invitrogen PA5-28382, 1:1000). ACTB (Sigma A5441, 1:10000) or GAPDH (Calbiochem CB1001, 1:10000) served as loading controls.

### Triple immunofluorescence

Mouse spinal cord immunohistochemistry was performed as previously described [81]. Here, the primary antibodies used were: ATXN2 (BD Transduction 611378, 1:50), PABPC1 (Abcam ab21060, 1:100); SQSTM1 (Santa Cruz sc25575, 1:50), VIM (Calbiochem LN6 clone V-9, 1:100). Secondary antibodies used were: Alexa Fluor 488 goat anti-mouse IgG A11029, Alexa Fluor 488 goat anti-rabbit IgG A11034, Alexa Fluor 565 rabbit anti-goat IgG A11079, Alexa Fluor 568 donkey anti-sheep IgG A21099, Alexa Fluor 568 goat anti-rabbit IgG A11036 (all Invitrogen, 1:1000).

### Coimmunoprecipitation and mass spectrometry

#### Sample preparation for mass spectrometry

Eluted IP-samples were incubated with DTT (5 mM final concentration) for 30 min at 55 °C. Samples were cooled down to room temperature and chloroacetamide was added to the reduced protein samples at a final concentration of 15 mM. Alkylation was carried out in the dark for 30 min at room temperature. The reaction was quenched with DTT (10 mM final concentration) for 15 min.

Samples were precipitated using four volumes of ice-cold methanol, one volume of chloroform and three volumes of ddH_2_O. After centrifugation at 14,000 g for 5 min at 4 °C, the upper aqueous phase was carefully aspirated, and three volumes of ice-cold methanol were added. Samples were mixed and centrifuged at 14,000 g for 5 min at 4 °C. Methanol was discarded and protein pellets were dried at room temperature.

Samples were re-suspended in 50 mM ammonium bicarbonate (ABC), 10% acetonitrile (ACN). Proteins were digested with 0.4 µg Trypsin (Promega, V5113) over night at 37 °C. Peptides were acidified using formic acid (FA) to a pH of 2-3 and dried in a vacuum centrifuge.

Peptide samples were purified using C18 (Octadecyl) resin material (3M Empore). C18 material was activated by incubation with methanol for 5 min and one wash with 50% ACN, 5% FA, followed by two additional washes with 3% ACN, 5% FA. Samples were re-suspended in 3% ACN, 5% FA and loaded to resin material. Peptides were washed with 3% ACN, 5% FA and eluted with 75% ACN, 5% FA. Samples were dried and re-suspended in 0.1% FA for LC-MS3.

#### Liquid chromatography mass spectrometry

Data acquisition was performed using centroid mode on an Orbitrap Fusion Lumos mass spectrometer and an easy-nLC 1200 nano HPLC system with a nanoFlex ion source (Thermo Fisher Scientific) as previously described [277, 278].

#### Mass spectrometry data analysis

Raw files of label-free single shots were analyzed using Proteome Discoverer (PD) v2.4 software (ThermoFisher Scientific). Database searches were run against trypsin-digested *Mus musculus* SwissProt (TaxID=10090, v2017-10-25) and FASTA files of common contaminants (’contaminants.fastà provided with MaxQuant) for quality control. Spectra were selected using default settings and database searches performed using SequestHT node. Static modifications were set as carbamidomethyl (+57.021464) at cysteine residues. Dynamic modifications were set as methionine oxidation (+15.995) and acetylation (+42.0106) at the N-terminus.

### Bioinformatics analyses

STRING web-server (https://string-db.org/) (version 11.0, last accessed on March 4, 2024) [279] was employed for the generation, visualization and analysis of protein-protein interaction networks, using the list of proteins with dysregulated phosphorylations identified in the *Atxn2*-CAG100-KIN 14-month-old mice spinal cord phosphorylation profiling, or the KIN cerebellar CoIP data as input. Connectivity scores and interactions between the proteins were extracted as tab delimited text files. The pathway enrichment analyses were conducted using different functional classification frameworks including Gene Ontology (GO), Kyoto Encyclopedia of Genes and Genomes (KEGG) Pathways, Reactome Pathways, and The Mammalian Phenotype Ontology (Monarch).

Disordered domains within ATXN2 and SQSTM1 were predicted using metapredict V2 (v2.4) (https://metapredict.net/, last accessed on February 29, 2024).

Phospho-sites and their regulation in mice and humans according to previously published literature were interrogated with help of the database at www.phosphositeplus.org (last accessed on 28 February 2024).

### Statistical analyses

Unpaired Student t-tests with Welch’s correction were used to establish comparisons for continuous variables between homozygous *Atxn2*-CAG100-KIN and WT animals. Bar charts depicting the mean and standard error of the mean (SEM) values were used for data visualization. All statistical analyses were conducted using GraphPad Prism software (version 8.4.2) (GraphPad Software Inc., USA). Significance was assumed at p<0.05 and highlighted with asterisks: Trend (T)<0.01; *p<0.05, **p<0.01, ***p<0.001, ****p<0.0001.

## Abbreviations

aka: also known as
ALS: amyotrophic lateral scleroosis
ATXN2: Ataxin-2
CAG: cytosine-adenine-guanine
CIN85: Cbl-interacting protein of 85 kDa
DLG4: discs large MAGUK scaffold protein 4
FC: fold change
FDR: false discovery rate
Fe-NTA: iron-nitrilotriacetic acid
FTD: fronto-temporal dementia
IMAC: immobilized metal affinity chromatography
KIN: KnockIn
KIR: Keap1-interacting region
KO: KnockOut
LIR: LC3-interaction region
LLPS: liquid-liquid phase separation
LSm: like-Sm domain in cytoplasmic proteins
LSm-AD: LSm-associated domain
MYO6: unconventional myosin VI
OPTN: optineurin
PABPC1: poly(A)-binding protein cytoplasmic 1
PAM2: poly(A)-binding-protein associated motif 2
polyQ: polyglutamine
PRM: proline-rich-motif
PSMD9: proteasome 19S regulator non-ATPase assembly chaperone P27
PSP: progressive supranuclear palsy
RPL21: ribosomal protein large subunit component 21
RTK: receptor tyrosine kinase
SAR: selective autophagy receptor
SCA2: spinocerebellar ataxia type 2
SH3: SRC-homology domain type 3
Sm: single-strand poly(C) binding domain in nuclear proteins
SPARCL1: secreted protein acidic and cysteine rich-like 1
STRING: search tool for the retrieval of interacting genes
TDP-43/TARDBP: TAR DNA-binding protein 43
TNIP1/ABIN1: TNFAIP3 interacting protein 1
UBQLN2: ubiquilin-2
WNK1: protein kinase with no lysine 1
WT: wildtype.

## Author Contributions

Conceptualization, S.G. and G.A.; methodology, J.C.-P., C.M. and S.G.; software, A.R.K., J.K., C.M.; validation, L.-E.A.-M., A.R.K., N.-E.S., J.C.-P., L.-M.B., J.K. and S.G.; formal analysis, C.M. and G.A.; investigation, L.-E.A.-M., A.R.K., M.E.B.; resources, N.-E.S., J.C.-P. and S.G.; data curation, C.M. and G.A.; writing—original draft preparation, L.E.A.-M., A.R.K., J.K. and G.A.; writing—review and editing, A.R.K., N.-E.S., J.K., C.M. and S.G.; visualization, L.-E.A.-M., A.R.K., J.C.-P., J.K. and G.A.; J.K., C.M., S.G. and G.A.; project administration, G.A.; funding acquisition, G.A. All authors have read and agreed to the published version of the manuscript.

## Funding

This research was funded by the Deutsche Forschungsgemeinschaft (grants AU96/11-1, 11-3,16-1 and 21-1 to GA, as well as grant ID 259130777 in the SFB1177 on Selective Autophagy to CM), and grant 403765277 for mass spectrometer to CM.

## Institutional Review Board Statement

The study was conducted in accordance with the Declaration of Helsinki. The analysis of human fibroblasts was approved by the Institutional Review Board (or Ethics Committee) of the Goethe University Frankfurt Medical School (protocol code 147/7, approved on June 26, 2007). The animal study protocol was approved by the Regierungspräsidium Darmstadt (protocol code V54-19c18-FK/1083, approved on March 27, 2017).

## Informed Consent Statement

Written informed consent was obtained from all subjects involved in the study.

## Data Availability Statement

The phosphoproteome and coimmunoprecipitation data are being deposited in the publically available repository PRIDE, and can be accessed with the identifiers (to be inserted), respectively.

## Acknowledgments

For technical assistance, we are grateful to Gabriele Koepf in the lab, and to the staff of the ZFE animal facility at the Goethe University in Frankfurt. For technical help and informatics advice regarding the phosphoproteome, we thank Dr. Matthew P. Stokes and his team at Cell Signaling Technologies INC., Danvers, MA, USA. We thank the Center for Functional Proteomics at the Goethe University in Frankfurt for mass spectrometry support.

## Conflicts of Interest

The authors declare no conflicts of interest. The funders had no role in the design of the study; in the collection, analyses, or interpretation of data; in the writing of the manuscript; or in the decision to publish the results.

## Appendix

**Figure S1.**
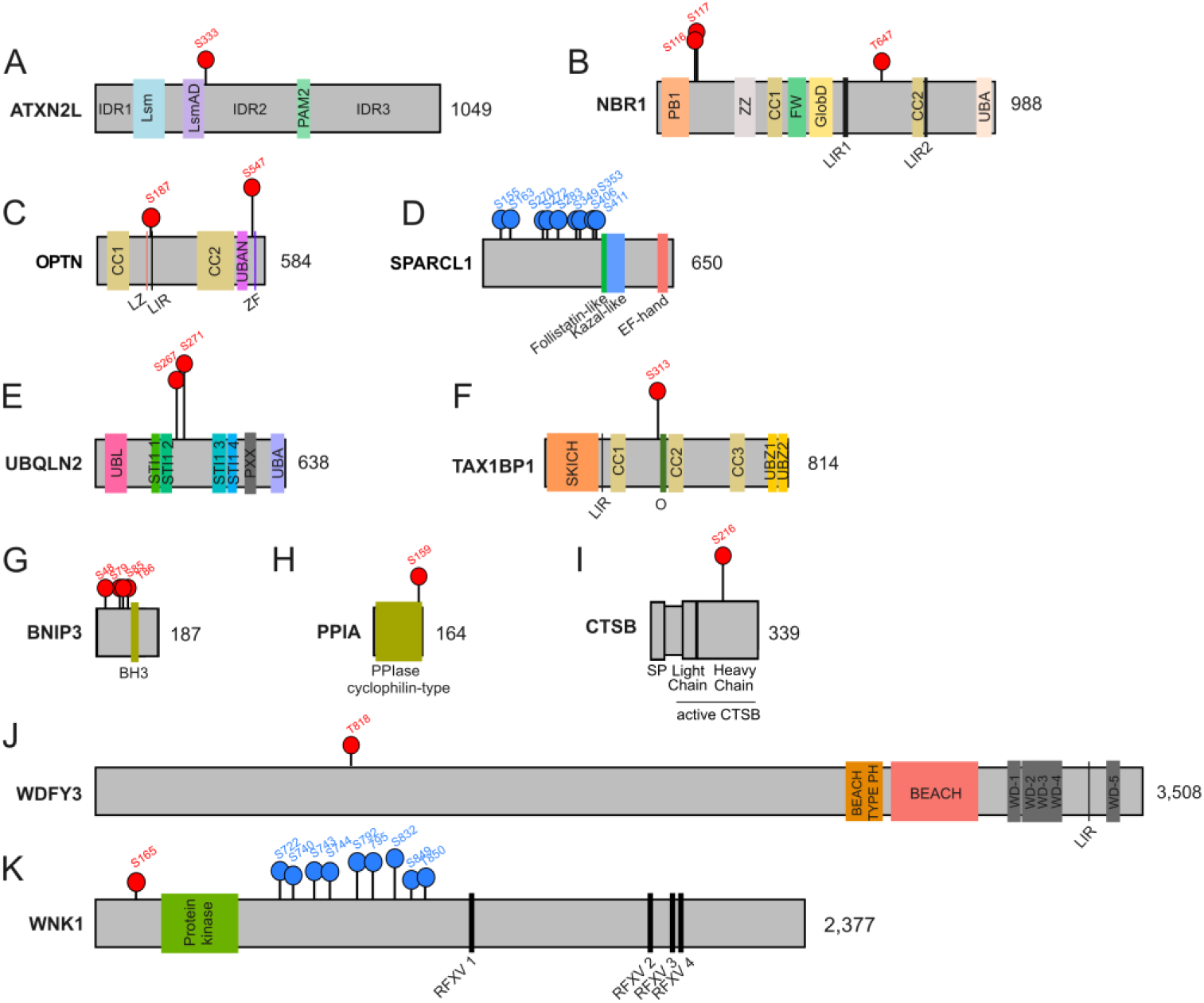
Schematic illustration of protein structures in ATXN2L, SPARCL1, and proteostasis/autophagy factors with their phosphorylation changes in aged KIN spinal cord.

**Figure S2.**
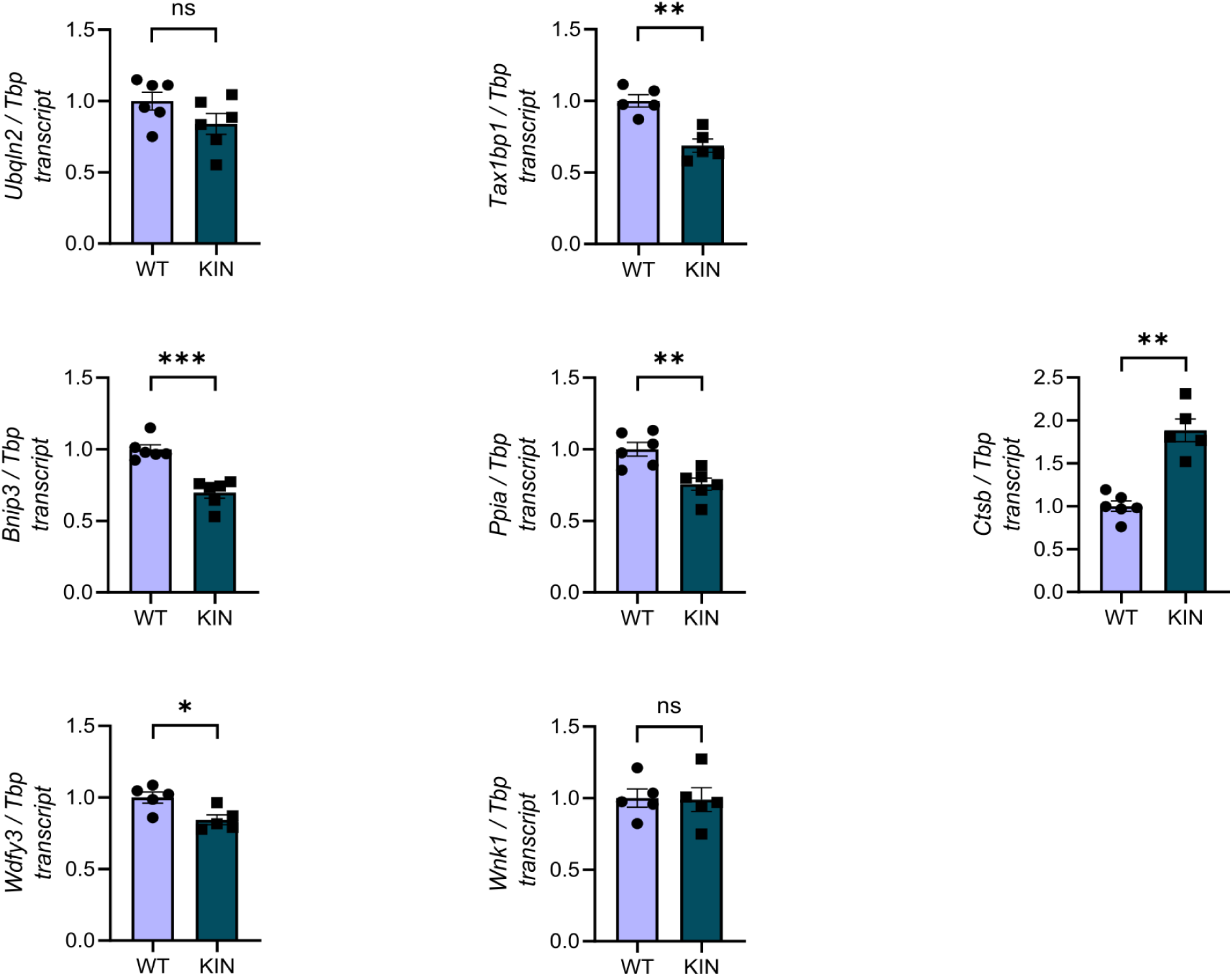
Quantitative reverse-transcriptase real-time PCR of *Ubqln2*, *Tax1bp1*, *Bnip3, Ppia, Ctsb, Wdfy3*, *Wnk1*, documenting the degree of altered mRNA expression in aged KIN spinal cord (N = 6 vs. 6).

**Figure S3.**
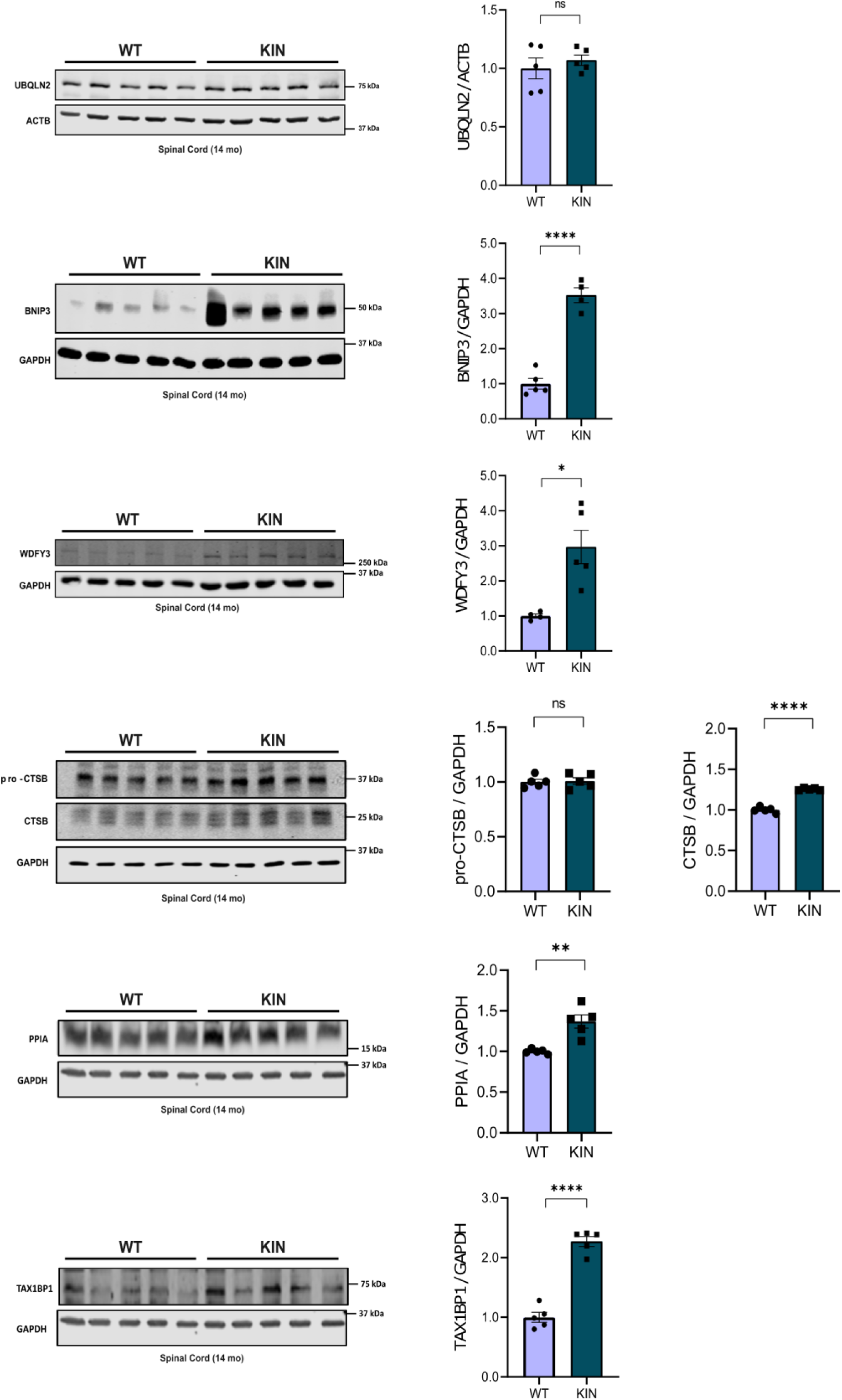
Quantitative immunoblots for UBQLN2, BNIP3, WDFY3 as autophagy regulators, CTSB as lysosomal enzyme, PPIA as ALS modulator, and TAX1BP1 as autophagosome maturation control, in end-stage spinal cord from our SCA2/ALS13 mouse model (KIN) versus wildtype (WT).

**Figure S4.**
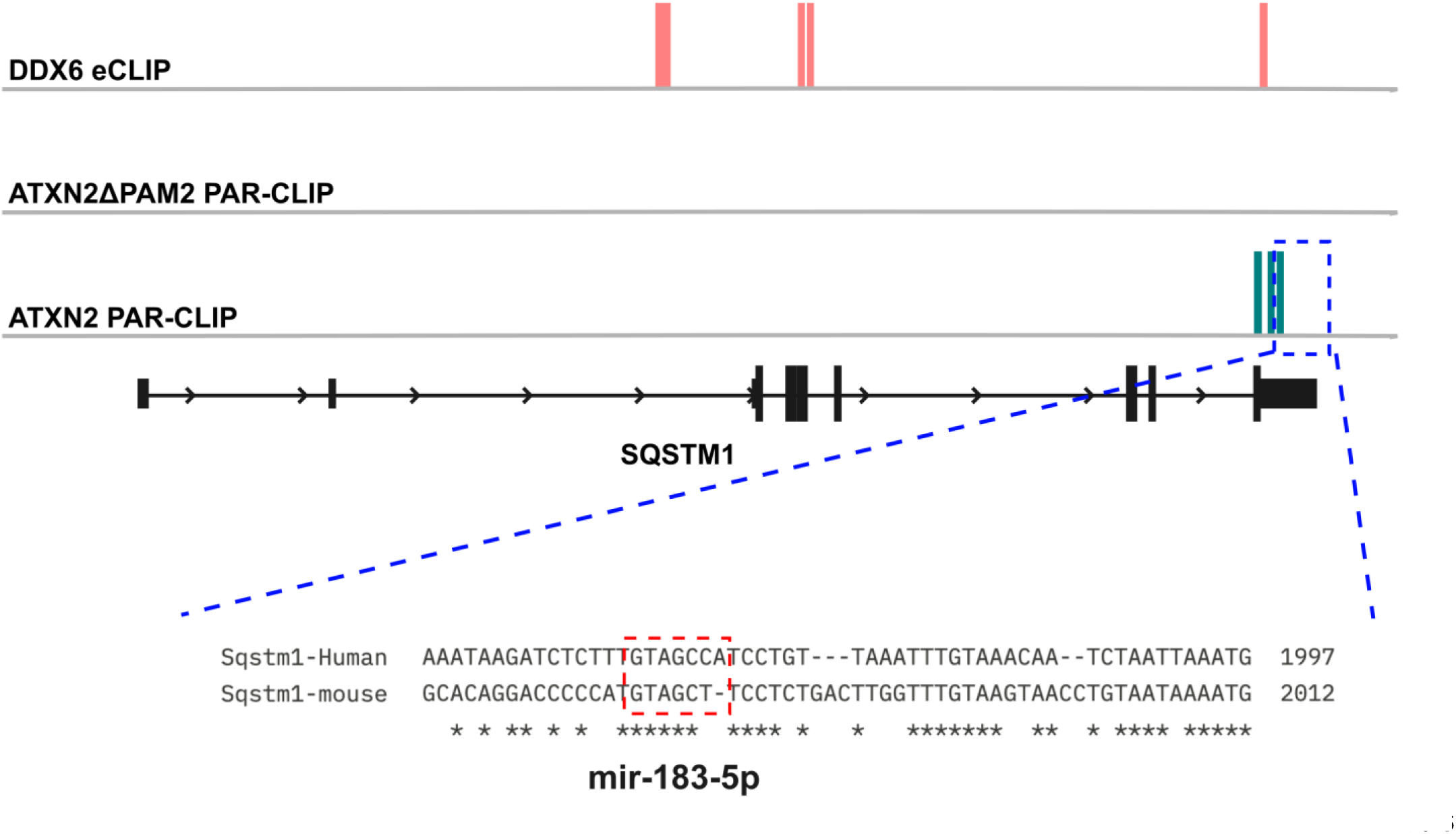
Schematic illustration of ATXN2 interaction areas with *Sqstm1* mRNA 3’UTR, the RNA helicase DDX6, and the microRNA-183-5p, according to publications that used eCLIP/PAR-CLIP technology. DDX6 data were taken from ENCODE database.

## Notes

### Competing Interest Statement

The authors have declared no competing interest.

## References

1. Pulst, S.M., et al., Moderate expansion of a normally biallelic trinucleotide repeat in spinocerebellar ataxia type 2. Nat Genet, 1996. 14(3): p. 269–76.

2. Sanpei, K., et al., Identification of the spinocerebellar ataxia type 2 gene using a direct identification of repeat expansion and cloning technique, DIRECT. Nat Genet, 1996. 14(3): p. 277–84.

3. Imbert, G., et al., Cloning of the gene for spinocerebellar ataxia 2 reveals a locus with high sensitivity to expanded CAG/glutamine repeats. Nat Genet, 1996. 14(3): p. 285–91.

4. Orozco Diaz, G., et al., Autosomal dominant cerebellar ataxia: clinical analysis of 263 patients from a homogeneous population in Holguin, Cuba. Neurology, 1990. 40(9): p. 1369–75.

5. Auburger, G.W., Spinocerebellar ataxia type 2. Handb Clin Neurol, 2012. 103: p. 423–36.

6. Almaguer-Mederos, L.E., et al., Estimation of survival in spinocerebellar ataxia type 2 Cuban patients. Clin Genet, 2013. 83(3): p. 293–4.

7. Rub, U., et al., Clinical features, neurogenetics and neuropathology of the polyglutamine spinocerebellar ataxias type 1, 2, 3, 6 and 7. Prog Neurobiol, 2013. 104: p. 38–66.

8. Almaguer-Mederos, L.E., et al., Body Mass Index Is Significantly Associated With Disease Severity in Spinocerebellar Ataxia Type 2 Patients. Mov Disord, 2021. 36(6): p. 1372–1380.

9. Elden, A.C., et al., Ataxin-2 intermediate-length polyglutamine expansions are associated with increased risk for ALS. Nature, 2010. 466(7310): p. 1069–75.

10. Lee, T., et al., Ataxin-2 intermediate-length polyglutamine expansions in European ALS patients. Hum Mol Genet, 2011. 20(9): p. 1697–700.

11. Gispert, S., et al., The modulation of Amyotrophic Lateral Sclerosis risk by ataxin-2 intermediate polyglutamine expansions is a specific effect. Neurobiol Dis, 2012. 45(1): p. 356–61.

12. Lahut, S., et al., ATXN2 and its neighbouring gene SH2B3 are associated with increased ALS risk in the Turkish population. PLoS One, 2012. 7(8): p. e42956.

13. Sproviero, W., et al., ATXN2 trinucleotide repeat length correlates with risk of ALS. Neurobiol Aging, 2017. 51: p. 178 e1–178 e9.

14. Chio, A., et al., Association of Copresence of Pathogenic Variants Related to Amyotrophic Lateral Sclerosis and Prognosis. Neurology, 2023. 101(1): p. e83–e93.

15. Fournier, C., et al., Interrupted CAG expansions in ATXN2 gene expand the genetic spectrum of frontotemporal dementias. Acta Neuropathol Commun, 2018. 6(1): p. 41.

16. Rubino, E., et al., ATXN2 intermediate repeat expansions influence the clinical phenotype in frontotemporal dementia. Neurobiol Aging, 2019. 73: p. 231 e7–231 e9.

17. Glass, J.D., et al., ATXN2 intermediate expansions in amyotrophic lateral sclerosis. Brain, 2022. 145(8): p. 2671–2676.

18. Ross, O.A., et al., Ataxin-2 repeat-length variation and neurodegeneration. Hum Mol Genet, 2011. 20(16): p. 3207–12.

19. Kim, Y.E., et al., SCA2 family presenting as typical Parkinson’s disease: 34 year follow up. Parkinsonism Relat Disord, 2017. 40: p. 69–72.

20. Kwong, L.K., et al., TDP-43 proteinopathy: the neuropathology underlying major forms of sporadic and familial frontotemporal lobar degeneration and motor neuron disease. Acta Neuropathol, 2007. 114(1): p. 63–70.

21. Liao, Y.Z., J. Ma, and J.Z. Dou, The Role of TDP-43 in Neurodegenerative Disease. Mol Neurobiol, 2022. 59(7): p. 4223–4241.

22. Highley, J.R., et al., Motor neurone disease/amyotrophic lateral sclerosis associated with intermediate-length CAG repeat expansions in Ataxin-2 does not have 1C2-positive polyglutamine inclusions. Neuropathol Appl Neurobiol, 2016. 42(4): p. 377–89.

23. Watanabe, R., et al., Intracellular dynamics of Ataxin-2 in the human brains with normal and frontotemporal lobar degeneration with TDP-43 inclusions. Acta Neuropathol Commun, 2020. 8(1): p. 176.

24. Koyano, S., et al., Parallel Appearance of Polyglutamine and Transactivation-Responsive DNA-Binding Protein 43 and Their Complementary Subcellular Localization in Brains of Patients With Spinocerebellar Ataxia Type 2. J Neuropathol Exp Neurol, 2022. 81(7): p. 535–544.

25. Wijegunawardana, D., et al., Ataxin-2 polyglutamine expansions aberrantly sequester TDP-43, drive ribonucleoprotein condensate transport dysfunction and suppress local translation. bioRxiv, 2023.

26. Becker, L.A., et al., Therapeutic reduction of ataxin-2 extends lifespan and reduces pathology in TDP-43 mice. Nature, 2017. 544(7650): p. 367–371.

27. Zeballos, C.M., et al., Mitigating a TDP-43 proteinopathy by targeting ataxin-2 using RNA-targeting CRISPR effector proteins. Nat Commun, 2023. 14(1): p. 6492.

28. Wijegunawardana, D., et al., Ataxin-2 polyglutamine expansions aberrantly sequester TDP-43 ribonucleoprotein condensates disrupting mRNA transport and local translation in neurons. Dev Cell, 2024.

29. Scoles, D.R., et al., Antisense oligonucleotide therapy for spinocerebellar ataxia type 2. Nature, 2017. 544(7650): p. 362–366.

30. Scoles, D.R., et al., A quantitative high-throughput screen identifies compounds that lower expression of the SCA2-and ALS-associated gene ATXN2. J Biol Chem, 2022. 298(8): p. 102228.

31. Afonso, I.T., et al., Mutant Ataxin-2 Expression in Aged Animals Aggravates Neuropathological Features Associated with Spinocerebellar Ataxia Type 2. Int J Mol Sci, 2022. 23(19).

32. Marcelo, A., et al., Autophagy in Spinocerebellar ataxia type 2, a dysregulated pathway, and a target for therapy. Cell Death Dis, 2021. 12(12): p. 1117.

33. Wardman, J.H., et al., Enhancement of Autophagy and Solubilization of Ataxin-2 Alleviate Apoptosis in Spinocerebellar Ataxia Type 2 Patient Cells. Cerebellum, 2020. 19(2): p. 165–181.

34. Lastres-Becker, I., et al., Insulin receptor and lipid metabolism pathology in ataxin-2 knock-out mice. Hum Mol Genet, 2008. 17(10): p. 1465–81.

35. Meierhofer, D., et al., Ataxin-2 (Atxn2)-Knock-Out Mice Show Branched Chain Amino Acids and Fatty Acids Pathway Alterations. Mol Cell Proteomics, 2016. 15(5): p. 1728–39.

36. Seidel, G., et al., Quantitative Global Proteomics of Yeast PBP1 Deletion Mutants and Their Stress Responses Identifies Glucose Metabolism, Mitochondrial, and Stress Granule Changes. J Proteome Res, 2017. 16(2): p. 504–515.

37. Key, J., et al., Mid-Gestation lethality of Atxn2l-Ablated Mice. Int J Mol Sci, 2020. 21(14).

38. Jimenez-Lopez, D. and P. Guzman, Insights into the evolution and domain structure of Ataxin-2 proteins across eukaryotes. BMC Res Notes, 2014. 7: p. 453.

39. Mangus, D.A., N. Amrani, and A. Jacobson, Pbp1p, a factor interacting with Saccharomyces cerevisiae poly(A)-binding protein, regulates polyadenylation. Mol Cell Biol, 1998. 18(12): p. 7383–96.

40. Kozlov, G., et al., Structure and function of the C-terminal PABC domain of human poly(A)-binding protein. Proc Natl Acad Sci U S A, 2001. 98(8): p. 4409–13.

41. Ciosk, R., M. DePalma, and J.R. Priess, ATX-2, the C. elegans ortholog of ataxin 2, functions in translational regulation in the germline. Development, 2004. 131(19): p. 4831–41.

42. Satterfield, T.F. and L.J. Pallanck, Ataxin-2 and its Drosophila homolog, ATX2, physically assemble with polyribosomes. Hum Mol Genet, 2006. 15(16): p. 2523–32.

43. van de Loo, S., et al., Ataxin-2 associates with rough endoplasmic reticulum. Exp Neurol, 2009. 215(1): p. 110–8.

44. Damrath, E., et al., ATXN2-CAG42 sequesters PABPC1 into insolubility and induces FBXW8 in cerebellum of old ataxic knock-in mice. PLoS Genet, 2012. 8(8): p. e1002920.

45. Lim, C. and R. Allada, ATAXIN-2 activates PERIOD translation to sustain circadian rhythms in Drosophila. Science, 2013. 340(6134): p. 875-9.

46. Zhang, Y., et al., A role for Drosophila ATX2 in activation of PER translation and circadian behavior. Science, 2013. 340(6134): p. 879-82.

47. Fittschen, M., et al., Genetic ablation of ataxin-2 increases several global translation factors in their transcript abundance but decreases translation rate. Neurogenetics, 2015. 16(3): p. 181–92.

48. Lastres-Becker, I., et al., Mammalian ataxin-2 modulates translation control at the pre-initiation complex via PI3K/mTOR and is induced by starvation. Biochim Biophys Acta, 2016. 1862(9): p. 1558–69.

49. Bakthavachalu, B., et al., RNP-Granule Assembly via Ataxin-2 Disordered Domains Is Required for Long-Term Memory and Neurodegeneration. Neuron, 2018. 98(4): p. 754–766 e4.

50. Inagaki, H., et al., Direct evidence that Ataxin-2 is a translational activator mediating cytoplasmic polyadenylation. J Biol Chem, 2020. 295(47): p. 15810–15825.

51. Nadimpalli, H.P., et al., Ataxin-2, Twenty-four, and Dicer-2 are components of a noncanonical cytoplasmic polyadenylation complex. Life Sci Alliance, 2022. 5(12).

52. Boeynaems, S., et al., Poly(A)-binding protein is an ataxin-2 chaperone that regulates biomolecular condensates. Mol Cell, 2023. 83(12): p. 2020–2034 e6.

53. Zhuang, Y., et al., Circadian clocks are modulated by compartmentalized oscillating translation. Cell, 2023. 186(15): p. 3245–3260 e23.

54. Albrecht, M. and T. Lengauer, Novel Sm-like proteins with long C-terminal tails and associated methyltransferases. FEBS Lett, 2004. 569(1-3): p. 18–26.

55. Albrecht, M., et al., Structural and functional analysis of ataxin-2 and ataxin-3. Eur J Biochem, 2004. 271(15): p. 3155–70.

56. Lekontseva, N.V., E.A. Stolboushkina, and A.D. Nikulin, Diversity of LSM Family Proteins: Similarities and Differences. Biochemistry (Mosc), 2021. 86(Suppl 1): p. S38–S49.

57. Yokoshi, M., et al., Direct binding of Ataxin-2 to distinct elements in 3’ UTRs promotes mRNA stability and protein expression. Mol Cell, 2014. 55(2): p. 186–98.

58. Nonhoff, U., et al., Ataxin-2 interacts with the DEAD/H-box RNA helicase DDX6 and interferes with P-bodies and stress granules. Mol Biol Cell, 2007. 18(4): p. 1385–96.

59. Lee, J., et al., LSM12 and ME31B/DDX6 Define Distinct Modes of Posttranscriptional Regulation by ATAXIN-2 Protein Complex in Drosophila Circadian Pacemaker Neurons. Mol Cell, 2017. 66(1): p. 129–140 e7.

60. Inagaki, H., N. Hosoda, and S.I. Hoshino, DDX6 is a positive regulator of Ataxin-2/PAPD4 cytoplasmic polyadenylation machinery. Biochem Biophys Res Commun, 2021. 553: p. 9–16.

61. Ralser, M., et al., An integrative approach to gain insights into the cellular function of human ataxin-2. J Mol Biol, 2005. 346(1): p. 203–14.

62. Swisher, K.D. and R. Parker, Localization to, and effects of Pbp1, Pbp4, Lsm12, Dhh1, and Pab1 on stress granules in Saccharomyces cerevisiae. PLoS One, 2010. 5(4): p. e10006.

63. Kaehler, C., et al., Ataxin-2-like is a regulator of stress granules and processing bodies. PLoS One, 2012. 7(11): p. e50134.

64. Wang, J.Y., et al., PolyQ-expanded ataxin-2 aggregation impairs cellular processing-body homeostasis via sequestering the RNA helicase DDX6. J Biol Chem, 2024. 300(7): p. 107413.

65. Nonis, D., et al., Ataxin-2 associates with the endocytosis complex and affects EGF receptor trafficking. Cell Signal, 2008. 20(10): p. 1725–39.

66. Ariumi, Y., et al., Hepatitis C virus hijacks P-body and stress granule components around lipid droplets. J Virol, 2011. 85(14): p. 6882–92.

67. Drost, J., et al., Ataxin-2 modulates the levels of Grb2 and SRC but not ras signaling. J Mol Neurosci, 2013. 51(1): p. 68–81.

68. Auburger, G., et al., Efficient Prevention of Neurodegenerative Diseases by Depletion of Starvation Response Factor Ataxin-2. Trends Neurosci, 2017. 40(8): p. 507–516.

69. Bar, D.Z., et al., Cell size and fat content of dietary-restricted Caenorhabditis elegans are regulated by ATX-2, an mTOR repressor. Proc Natl Acad Sci U S A, 2016. 113(32): p. E4620–9.

70. Kato, M., et al., Redox State Controls Phase Separation of the Yeast Ataxin-2 Protein via Reversible Oxidation of Its Methionine-Rich Low-Complexity Domain. Cell, 2019. 177(3): p. 711–721 e8.

71. Yang, Y.S., et al., Yeast Ataxin-2 Forms an Intracellular Condensate Required for the Inhibition of TORC1 Signaling during Respiratory Growth. Cell, 2019. 177(3): p. 697–710 e17.

72. DeMille, D., et al., PAS kinase is activated by direct SNF1-dependent phosphorylation and mediates inhibition of TORC1 through the phosphorylation and activation of Pbp1. Mol Biol Cell, 2015. 26(3): p. 569–82.

73. Liu, Y.J., et al., Ataxin-2 sequesters Raptor into aggregates and impairs cellular mTORC1 signaling. FEBS J, 2024. 291(8): p. 1795–1812.

74. Prouteau, M. and R. Loewith, TOR Signaling Is Going through a Phase. Cell Metab, 2019. 29(5): p. 1019–1021.

75. Satterfield, T.F., S.M. Jackson, and L.J. Pallanck, A Drosophila homolog of the polyglutamine disease gene SCA2 is a dosage-sensitive regulator of actin filament formation. Genetics, 2002. 162(4): p. 1687–702.

76. Sen, N.E. and G. Auburger, Modeling amyotrophic lateral sclerosis through Ataxin-2 pathology, in Handbook of Animal Models in Neurological Disorders, V.R. Preedy, C. R. Martin, and V.B. Patel, Editors. 2023, Acadmic Press: Cambridge, MA. p. Pages 95–106.

77. Sen, N.E., et al., Generation of an Atxn2-CAG100 knock-in mouse reveals N-acetylaspartate production deficit due to early Nat8l dysregulation. Neurobiol Dis, 2019. 132: p. 104559.

78. Bux, J., et al., TR-FRET-Based Immunoassay to Measure Ataxin-2 as a Target Engagement Marker in Spinocerebellar Ataxia Type 2. Mol Neurobiol, 2023. 60(6): p. 3553–3567.

79. Arsovic, A., et al., Mouse Ataxin-2 Expansion Downregulates CamKII and Other Calcium Signaling Factors, Impairing Granule-Purkinje Neuron Synaptic Strength. Int J Mol Sci, 2020. 21(18).

80. Sen, N.E., et al., In Human and Mouse Spino-Cerebellar Tissue, Ataxin-2 Expansion Affects Ceramide-Sphingomyelin Metabolism. Int J Mol Sci, 2019. 20(23).

81. Canet-Pons, J., et al., Atxn2-CAG100-KnockIn mouse spinal cord shows progressive TDP43 pathology associated with cholesterol biosynthesis suppression. Neurobiol Dis, 2021. 152: p. 105289.

82. Ragothaman, M., et al., Complex phenotypes in an Indian family with homozygous SCA2 mutations. Ann Neurol, 2004. 55(1): p. 130–3.

83. Hoche, F., et al., Spinocerebellar ataxia type 2 (SCA2): identification of early brain degeneration in one monozygous twin in the initial disease stage. Cerebellum, 2011. 10(2): p. 245–53.

84. Tojima, M., et al., Homozygous 31 trinucleotide repeats in the SCA2 allele are pathogenic for cerebellar ataxia. Neurol Genet, 2018. 4(6): p. e283.

85. Uversky, V.N., The roles of intrinsic disorder-based liquid-liquid phase transitions in the “Dr. Jekyll-Mr. Hyde” behavior of proteins involved in amyotrophic lateral sclerosis and frontotemporal lobar degeneration. Autophagy, 2017. 13(12): p. 2115–2162.

86. Li, P., et al., Phase transitions in the assembly of multivalent signalling proteins. Nature, 2012. 483(7389): p. 336-40.

87. Amaya, J., V.H. Ryan, and N.L. Fawzi, The SH3 domain of Fyn kinase interacts with and induces liquid-liquid phase separation of the low-complexity domain of hnRNPA2. J Biol Chem, 2018. 293(51): p. 19522–19531.

88. Ghosh, A., K. Mazarakos, and H.X. Zhou, Three archetypical classes of macromolecular regulators of protein liquid-liquid phase separation. Proc Natl Acad Sci U S A, 2019. 116(39): p. 19474–19483.

89. Pankivskyi, S., et al., ITSN1 regulates SAM68 solubility through SH3 domain interactions with SAM68 proline-rich motifs. Cell Mol Life Sci, 2021. 78(4): p. 1745–1763.

90. Martinez-Marmol, R., et al., Fyn nanoclustering requires switching to an open conformation and is enhanced by FTLD-Tau biomolecular condensates. Mol Psychiatry, 2023. 28(2): p. 946–962.

91. Lopez-Palacios, T.P. and J.L. Andersen, Kinase regulation by liquid-liquid phase separation. Trends Cell Biol, 2023. 33(8): p. 649–666.

92. Maier, J., et al., Quantitative description of the phase-separation behavior of the multivalent SLP65-CIN85 complex. PNAS Nexus, 2024. 3(3): p. pgae079.

93. Kazemein Jasemi, N.S., et al., Functional Classification and Interaction Selectivity Landscape of the Human SH3 Domain Superfamily. Cells, 2024. 13(2).

94. Umemori, H., et al., Initial events of myelination involve Fyn tyrosine kinase signalling. Nature, 1994. 367(6463): p. 572-6.

95. Yamada, E., et al., Mouse skeletal muscle fiber-type-specific macroautophagy and muscle wasting are regulated by a Fyn/STAT3/Vps34 signaling pathway. Cell Rep, 2012. 1(5): p. 557–69.

96. Yang, M., et al., A C9ORF72/SMCR8-containing complex regulates ULK1 and plays a dual role in autophagy. Sci Adv, 2016. 2(9): p. e1601167.

97. Ciura, S., et al., The most prevalent genetic cause of ALS-FTD, C9orf72 synergizes the toxicity of ATXN2 intermediate polyglutamine repeats through the autophagy pathway. Autophagy, 2016. 12(8): p. 1406–8.

98. Sellier, C., et al., Loss of C9ORF72 impairs autophagy and synergizes with polyQ Ataxin-2 to induce motor neuron dysfunction and cell death. EMBO J, 2016. 35(12): p. 1276–97.

99. Ke, P.Y., The Multifaceted Roles of Autophagy in Flavivirus-Host Interactions. Int J Mol Sci, 2018. 19(12).

100. Conway, O., et al., Selective Autophagy Receptors in Neuronal Health and Disease. J Mol Biol, 2020. 432(8): p. 2483–2509.

101. Yang, C.S., et al., Autophagy protein Rubicon mediates phagocytic NADPH oxidase activation in response to microbial infection or TLR stimulation. Cell Host Microbe, 2012. 11(3): p. 264–76.

102. Cipriano, A., et al., NADPH Oxidases: From Molecular Mechanisms to Current Inhibitors. J Med Chem, 2023. 66(17): p. 11632–11655.

103. Gallolu Kankanamalage, S., et al., Multistep regulation of autophagy by WNK1. Proc Natl Acad Sci U S A, 2016. 113(50): p. 14342–14347.

104. McCormick, J.A. and D.H. Ellison, The WNKs: atypical protein kinases with pleiotropic actions. Physiol Rev, 2011. 91(1): p. 177–219.

105. Yamamoto, Y., et al., NEK9 regulates primary cilia formation by acting as a selective autophagy adaptor for MYH9/myosin IIA. Nat Commun, 2021. 12(1): p. 3292.

106. Cha-Molstad, H., et al., p62/SQSTM1/Sequestosome-1 is an N-recognin of the N-end rule pathway which modulates autophagosome biogenesis. Nat Commun, 2017. 8(1): p. 102.

107. Cha-Molstad, H., et al., Regulation of autophagic proteolysis by the N-recognin SQSTM1/p62 of the N-end rule pathway. Autophagy, 2018. 14(2): p. 359–361.

108. Ji, C.H., et al., The N-Degron Pathway Mediates ER-phagy. Mol Cell, 2019. 75(5): p. 1058–1072 e9.

109. Bonnet, L.V., et al., Arginyltransferase 1 modulates p62-driven autophagy via mTORC1/AMPk signaling. Cell Commun Signal, 2024. 22(1): p. 87.

110. Varland, S., J. Vandekerckhove, and A. Drazic, Actin Post-translational Modifications: The Cinderella of Cytoskeletal Control. Trends Biochem Sci, 2019. 44(6): p. 502–516.

111. Chin, S.M., et al., N-terminal acetylation and arginylation of actin determines the architecture and assembly rate of linear and branched actin networks. J Biol Chem, 2022. 298(11): p. 102518.

112. Sahlender, D.A., et al., Optineurin links myosin VI to the Golgi complex and is involved in Golgi organization and exocytosis. J Cell Biol, 2005. 169(2): p. 285–95.

113. Ryan, T.A. and D.A. Tumbarello, Optineurin: A Coordinator of Membrane-Associated Cargo Trafficking and Autophagy. Front Immunol, 2018. 9: p. 1024.

114. Zhao, S., et al., Fundamental roles of the Optineurin gene in the molecular pathology of Amyotrophic Lateral Sclerosis. Front Neurosci, 2023. 17: p. 1319706.

115. Laplantine, E., et al., NEMO specifically recognizes K63-linked poly-ubiquitin chains through a new bipartite ubiquitin-binding domain. EMBO J, 2009. 28(19): p. 2885–95.

116. Gan, K.J. and T.C. Sudhof, SPARCL1 Promotes Excitatory But Not Inhibitory Synapse Formation and Function Independent of Neurexins and Neuroligins. J Neurosci, 2020. 40(42): p. 8088–8102.

117. Chen, S., et al., Regulation of SPARC family proteins in disorders of the central nervous system. Brain Res Bull, 2020. 163: p. 178–189.

118. Shimizu, H., et al., Sporadic ALS with compound heterozygous mutations in the SQSTM1 gene. Acta Neuropathol, 2013. 126(3): p. 453–9.

119. Soo, K.Y., et al., ALS-associated mutant FUS inhibits macroautophagy which is restored by overexpression of Rab1. Cell Death Discov, 2015. 1: p. 15030.

120. Madill, M., et al., Amyotrophic lateral sclerosis patient iPSC-derived astrocytes impair autophagy via non-cell autonomous mechanisms. Mol Brain, 2017. 10(1): p. 22.

121. Chitiprolu, M., et al., A complex of C9ORF72 and p62 uses arginine methylation to eliminate stress granules by autophagy. Nat Commun, 2018. 9(1): p. 2794.

122. Davidson, J.M., R.S. Chung, and A. Lee, The converging roles of sequestosome-1/p62 in the molecular pathways of amyotrophic lateral sclerosis (ALS) and frontotemporal dementia (FTD). Neurobiol Dis, 2022. 166: p. 105653.

123. Millecamps, S., et al., Screening of OPTN in French familial amyotrophic lateral sclerosis. Neurobiol Aging, 2011. 32(3): p. 557 e11-3.

124. van Blitterswijk, M., et al., Novel optineurin mutations in sporadic amyotrophic lateral sclerosis patients. Neurobiol Aging, 2012. 33(5): p. 1016 e1-7.

125. Kamada, M., et al., Clinicopathologic features of autosomal recessive amyotrophic lateral sclerosis associated with optineurin mutation. Neuropathology, 2014. 34(1): p. 64–70.

126. Pottier, C., et al., Whole-genome sequencing reveals important role for TBK1 and OPTN mutations in frontotemporal lobar degeneration without motor neuron disease. Acta Neuropathol, 2015. 130(1): p. 77–92.

127. Gotkine, M., et al., A recessive S174X mutation in Optineurin causes amyotrophic lateral sclerosis through a loss of function via allele-specific nonsense-mediated decay. Neurobiol Aging, 2021. 106: p. 351 e1–351 e6.

128. Prtenjaca, N., et al., Optineurin Deficiency and Insufficiency Lead to Higher Microglial TDP-43 Protein Levels. Int J Mol Sci, 2022. 23(12).

129. Li, C., et al., MicroRNA-183-5p is stress-inducible and protects neurons against cell death in amyotrophic lateral sclerosis. J Cell Mol Med, 2020. 24(15): p. 8614–8622.

130. Kim, H.C., et al., MicroRNA-183-5p regulates TAR DNA-binding protein 43 neurotoxicity via SQSTM1/p62 in amyotrophic lateral sclerosis. J Neurochem, 2023. 164(5): p. 643–657.

131. Klionsky, D.J., et al., Does bafilomycin A1 block the fusion of autophagosomes with lysosomes? Autophagy, 2008. 4(7): p. 849–50.

132. McEwen, E., et al., Heme-regulated inhibitor kinase-mediated phosphorylation of eukaryotic translation initiation factor 2 inhibits translation, induces stress granule formation, and mediates survival upon arsenite exposure. J Biol Chem, 2005. 280(17): p. 16925–33.

133. Paul, S., et al., Staufen Impairs Autophagy in Neurodegeneration. Ann Neurol, 2023. 93(2): p. 398–416.

134. Shi, J., et al., Dominant-negative function of the C-terminal fragments of NBR1 and SQSTM1 generated during enteroviral infection. Cell Death Differ, 2014. 21(9): p. 1432–41.

135. Sanchez-Martin, P., et al., NBR1-mediated p62-liquid droplets enhance the Keap1-Nrf2 system. EMBO Rep, 2020. 21(3): p. e48902.

136. Kozlov, G., et al., Structural basis of binding of P-body-associated proteins GW182 and ataxin-2 by the Mlle domain of poly(A)-binding protein. J Biol Chem, 2010. 285(18): p. 13599–606.

137. Zhang, C., et al., p38delta MAPK regulates aggresome biogenesis by phosphorylating SQSTM1 in response to proteasomal stress. J Cell Sci, 2018. 131(14).

138. Lauranzano, E., et al., Peptidylprolyl isomerase A governs TARDBP function and assembly in heterogeneous nuclear ribonucleoprotein complexes. Brain, 2015. 138(Pt 4): p. 974–91.

139. Pasetto, L., et al., Defective cyclophilin A induces TDP-43 proteinopathy: implications for amyotrophic lateral sclerosis and frontotemporal dementia. Brain, 2021. 144(12): p. 3710–3726.

140. Pasetto, L., et al., Targeting Extracellular Cyclophilin A Reduces Neuroinflammation and Extends Survival in a Mouse Model of Amyotrophic Lateral Sclerosis. J Neurosci, 2017. 37(6): p. 1413–1427.

141. Kimura, Y., K. Irie, and K. Irie, Pbp1 is involved in Ccr4- and Khd1-mediated regulation of cell growth through association with ribosomal proteins Rpl12a and Rpl12b. Eukaryot Cell, 2013. 12(6): p. 864–74.

142. Wawiorka, L., et al., Functional analysis of the uL11 protein impact on translational machinery. Cell Cycle, 2016. 15(8): p. 1060–72.

143. Imami, K., et al., Phosphorylation of the Ribosomal Protein RPL12/uL11 Affects Translation during Mitosis. Mol Cell, 2018. 72(1): p. 84–98 e9.

144. Molon, M., et al., Ribosomal Protein uL11 as a Regulator of Metabolic Circuits Related to Aging and Cell Cycle. Cells, 2020. 9(7).

145. Inglis, A.J., et al., Activation of GCN2 by the ribosomal P-stalk. Proc Natl Acad Sci U S A, 2019. 116(11): p. 4946–4954.

146. Masson, G.R., Towards a model of GCN2 activation. Biochem Soc Trans, 2019. 47(5): p. 1481–1488.

147. Harding, H.P., et al., The ribosomal P-stalk couples amino acid starvation to GCN2 activation in mammalian cells. Elife, 2019. 8.

148. Gupta, R. and A.G. Hinnebusch, Differential requirements for P stalk components in activating yeast protein kinase Gcn2 by stalled ribosomes during stress. Proc Natl Acad Sci U S A, 2023. 120(16): p. e2300521120.

149. Bou-Nader, C., et al., Gcn2 structurally mimics and functionally repurposes the HisRS enzyme for the integrated stress response. Proc Natl Acad Sci U S A, 2024. 121(35): p. e2409628121.

150. Costa-Mattioli, M., et al., Translational control of hippocampal synaptic plasticity and memory by the eIF2alpha kinase GCN2. Nature, 2005. 436(7054): p. 1166-73.

151. Roffe, M., et al., IMPACT is a developmentally regulated protein in neurons that opposes the eukaryotic initiation factor 2alpha kinase GCN2 in the modulation of neurite outgrowth. J Biol Chem, 2013. 288(15): p. 10860–9.

152. Shen, W.C., et al., Mutations in the ubiquitin-binding domain of OPTN/optineurin interfere with autophagy-mediated degradation of misfolded proteins by a dominant-negative mechanism. Autophagy, 2015. 11(4): p. 685–700.

153. Masters, T.A., et al., MYO6 Regulates Spatial Organization of Signaling Endosomes Driving AKT Activation and Actin Dynamics. Cell Rep, 2017. 19(10): p. 2088–2101.

154. Kruppa, A.J., et al., Myosin VI-Dependent Actin Cages Encapsulate Parkin-Positive Damaged Mitochondria. Dev Cell, 2018. 44(4): p. 484–499 e6.

155. de Jonge, J.J., et al., The MYO6 interactome: selective motor-cargo complexes for diverse cellular processes. FEBS Lett, 2019. 593(13): p. 1494–1507.

156. Qiu, Y., et al., Emerging views of OPTN (optineurin) function in the autophagic process associated with disease. Autophagy, 2022. 18(1): p. 73–85.

157. Mangiarini, L., et al., Exon 1 of the HD gene with an expanded CAG repeat is sufficient to cause a progressive neurological phenotype in transgenic mice. Cell, 1996. 87(3): p. 493–506.

158. Sittler, A., et al., SH3GL3 associates with the Huntingtin exon 1 protein and promotes the formation of polyglncontaining protein aggregates. Mol Cell, 1998. 2(4): p. 427–36.

159. Murphy, K.P., et al., Abnormal synaptic plasticity and impaired spatial cognition in mice transgenic for exon 1 of the human Huntington’s disease mutation. J Neurosci, 2000. 20(13): p. 5115–23.

160. Song, C., et al., Expression of polyglutamine-expanded huntingtin induces tyrosine phosphorylation of N-methyl-D-aspartate receptors. J Biol Chem, 2003. 278(35): p. 33364–9.

161. Qin, Z.H., et al., Huntingtin bodies sequester vesicle-associated proteins by a polyproline-dependent interaction. J Neurosci, 2004. 24(1): p. 269–81.

162. Ralser, M., et al., Ataxin-2 and huntingtin interact with endophilin-A complexes to function in plastin-associated pathways. Hum Mol Genet, 2005. 14(19): p. 2893–909.

163. Dehay, B. and A. Bertolotti, Critical role of the proline-rich region in Huntingtin for aggregation and cytotoxicity in yeast. J Biol Chem, 2006. 281(47): p. 35608–15.

164. Gao, Y.G., et al., Structural insights into the specific binding of huntingtin proline-rich region with the SH3 and WW domains. Structure, 2006. 14(12): p. 1755–65.

165. Southwell, A.L., et al., Intrabodies binding the proline-rich domains of mutant huntingtin increase its turnover and reduce neurotoxicity. J Neurosci, 2008. 28(36): p. 9013–20.

166. Landles, C., et al., Proteolysis of mutant huntingtin produces an exon 1 fragment that accumulates as an aggregated protein in neuronal nuclei in Huntington disease. J Biol Chem, 2010. 285(12): p. 8808–23.

167. Sathasivam, K., et al., Aberrant splicing of HTT generates the pathogenic exon 1 protein in Huntington disease. Proc Natl Acad Sci U S A, 2013. 110(6): p. 2366–70.

168. Crick, S.L., et al., Unmasking the roles of N- and C-terminal flanking sequences from exon 1 of huntingtin as modulators of polyglutamine aggregation. Proc Natl Acad Sci U S A, 2013. 110(50): p. 20075–80.

169. Baksi, S., S. Basu, and D. Mukhopadhyay, Mutant huntingtin replaces Gab1 and interacts with C-terminal SH3 domain of growth factor receptor binding protein 2 (Grb2). Neurosci Res, 2014. 87: p. 77–83.

170. Neueder, A., et al., The pathogenic exon 1 HTT protein is produced by incomplete splicing in Huntington’s disease patients. Sci Rep, 2017. 7(1): p. 1307.

171. Peskett, T.R., et al., A Liquid to Solid Phase Transition Underlying Pathological Huntingtin Exon1 Aggregation. Mol Cell, 2018. 70(4): p. 588–601 e6.

172. Franich, N.R., et al., Phenotype onset in Huntington’s disease knock-in mice is correlated with the incomplete splicing of the mutant huntingtin gene. J Neurosci Res, 2019. 97(12): p. 1590–1605.

173. Ceccon, A., V. Tugarinov, and G.M. Clore, Quantitative Exchange NMR-Based Analysis of Huntingtin-SH3 Interactions Suggests an Allosteric Mechanism of Inhibition of Huntingtin Aggregation. J Am Chem Soc, 2021. 143(25): p. 9672–9681.

174. Fao, L., et al., Restoration of c-Src/Fyn Proteins Rescues Mitochondrial Dysfunction in Huntington’s Disease. Antioxid Redox Signal, 2023. 38(1-3): p. 95–114.

175. Fao, L., et al., Restored Fyn Levels in Huntington’s Disease Contributes to Enhanced Synaptic GluN2B-Composed NMDA Receptors and CREB Activity. Cells, 2022. 11(19).

176. Brady, S.T., et al., Toxic effects of mutant huntingtin in axons are mediated by its proline-rich domain. Brain, 2024. 147(6): p. 2098–2113.

177. Hoschek, F., et al., Huntingtin HTT1a is generated in a CAG repeat-length-dependent manner in human tissues. Mol Med, 2024. 30(1): p. 36.

178. Sogorb-Gonzalez, M., et al., Exon 1-targeting miRNA reduces the pathogenic exon 1 HTT protein in Huntington disease models. Brain, 2024.

179. Chitre, M. and P. Emery, ATXN2 is a target of N-terminal proteolysis. PLoS One, 2023. 18(12): p. e0296085.

180. Wang, K.Z., et al., TRAF6 activation of PI 3-kinase-dependent cytoskeletal changes is cooperative with Ras and is mediated by an interaction with cytoplasmic Src. J Cell Sci, 2006. 119(Pt 8): p. 1579–91.

181. Scoles, D.R., et al., ETS1 regulates the expression of ATXN2. Hum Mol Genet, 2012. 21(23): p. 5048–65.

182. Monteiro, P. and G. Feng, SHANK proteins: roles at the synapse and in autism spectrum disorder. Nat Rev Neurosci, 2017. 18(3): p. 147–157.

183. Guergueltcheva, V., et al., Autosomal-recessive congenital cerebellar ataxia is caused by mutations in metabotropic glutamate receptor 1. Am J Hum Genet, 2012. 91(3): p. 553–64.

184. Watson, L.M., et al., Dominant Mutations in GRM1 Cause Spinocerebellar Ataxia Type 44. Am J Hum Genet, 2017. 101(3): p. 451–458.

185. Collins, M.A., et al., Label-Free LC-MS/MS Proteomic Analysis of Cerebrospinal Fluid Identifies Protein/Pathway Alterations and Candidate Biomarkers for Amyotrophic Lateral Sclerosis. J Proteome Res, 2015. 14(11): p. 4486–501.

186. Mongredien, R., et al., Cartography of hevin-expressing cells in the adult brain reveals prominent expression in astrocytes and parvalbumin neurons. Brain Struct Funct, 2019. 224(3): p. 1219–1244.

187. Kramer-Albers, E.M. and R. White, From axon-glial signalling to myelination: the integrating role of oligodendroglial Fyn kinase. Cell Mol Life Sci, 2011. 68(12): p. 2003–12.

188. Kliche, S., et al., The ADAP/SKAP55 signaling module regulates T-cell receptor-mediated integrin activation through plasma membrane targeting of Rap1. Mol Cell Biol, 2006. 26(19): p. 7130–44.

189. Ganesan, A., P. Muralidharan, and L.N. Ramya, The Fulcrum of Demyelination in Multiple Sclerosis. Curr Protein Pept Sci, 2023. 24(7): p. 579–588.

190. Wilkinson, B. and M.P. Coba, Molecular architecture of postsynaptic Interactomes. Cell Signal, 2020. 76: p. 109782.

191. Piccini, A. and R. Malinow, Critical postsynaptic density 95/disc large/zonula occludens-1 interactions by glutamate receptor 1 (GluR1) and GluR2 required at different subcellular sites. J Neurosci, 2002. 22(13): p. 5387–92.

192. Lua, B.L. and B.C. Low, Cortactin phosphorylation as a switch for actin cytoskeletal network and cell dynamics control. FEBS Lett, 2005. 579(3): p. 577–85.

193. Clements, C.M., et al., The Structural Dynamics, Complexity of Interactions, and Functions in Cancer of Multi-SAM Containing Proteins. Cancers (Basel), 2023. 15(11).

194. Liu, X., et al., PRG-1 Regulates Synaptic Plasticity via Intracellular PP2A/beta1-Integrin Signaling. Dev Cell, 2016. 38(3): p. 275–90.

195. Lin, W.H., et al., Vasodilator-stimulated phosphoprotein (VASP) induces actin assembly in dendritic spines to promote their development and potentiate synaptic strength. J Biol Chem, 2010. 285(46): p. 36010–20.

196. Schiapparelli, L.M., et al., Activity-Induced Cortical Glutamatergic Neuron Nascent Proteins. J Neurosci, 2022. 42(42): p. 7900–7920.

197. Nobuhisa, I., et al., Activation of the superoxide-producing phagocyte NADPH oxidase requires co-operation between the tandem SH3 domains of p47phox in recognition of a polyproline type II helix and an adjacent alpha-helix of p22phox. Biochem J, 2006. 396(1): p. 183–92.

198. Hayashi, T., et al., PX-RICS, a novel splicing variant of RICS, is a main isoform expressed during neural development. Genes Cells, 2007. 12(8): p. 929–39.

199. Liu, X., et al., A novel AHI-1-BCR-ABL-DNM2 complex regulates leukemic properties of primitive CML cells through enhanced cellular endocytosis and ROS-mediated autophagy. Leukemia, 2017. 31(11): p. 2376–2387.

200. Gallolu Kankanamalage, S., et al., WNK1 is an unexpected autophagy inhibitor. Autophagy, 2017. 13(5): p. 969–970.

201. van Vliet, A.R., et al., ATG9A and ATG2A form a heteromeric complex essential for autophagosome formation. Mol Cell, 2022. 82(22): p. 4324–4339 e8.

202. Nozawa, T., et al., Rab41-mediated ESCRT machinery repairs membrane rupture by a bacterial toxin in xenophagy. Nat Commun, 2023. 14(1): p. 6230.

203. Imamura, K., et al., The Src/c-Abl pathway is a potential therapeutic target in amyotrophic lateral sclerosis. Sci Transl Med, 2017. 9(391).

204. Chen, R., et al., Downregulation of ASPP2 improves hepatocellular carcinoma cells survival via promoting BECN1-dependent autophagy initiation. Cell Death Dis, 2016. 7(12): p. e2512.

205. Pang, Y., et al., TP53BP2 decreases cell proliferation and induces autophagy in neuroblastoma cell lines. Oncol Lett, 2019. 17(6): p. 4976–4984.

206. Kim, J.Y., et al., Dissection of TBK1 signaling via phosphoproteomics in lung cancer cells. Proc Natl Acad Sci U S A, 2013. 110(30): p. 12414–9.

207. Freischmidt, A., et al., Haploinsufficiency of TBK1 causes familial ALS and fronto-temporal dementia. Nat Neurosci, 2015. 18(5): p. 631–6.

208. Heath, R.J., et al., RNF166 Determines Recruitment of Adaptor Proteins during Antibacterial Autophagy. Cell Rep, 2016. 17(9): p. 2183–2194.

209. Ma, S., I.Y. Attarwala, and X.Q. Xie, SQSTM1/p62: A Potential Target for Neurodegenerative Disease. ACS Chem Neurosci, 2019. 10(5): p. 2094–2114.

210. Nakazawa, S., et al., Linear ubiquitination is involved in the pathogenesis of optineurin-associated amyotrophic lateral sclerosis. Nat Commun, 2016. 7: p. 12547.

211. Richter, B., et al., Phosphorylation of OPTN by TBK1 enhances its binding to Ub chains and promotes selective autophagy of damaged mitochondria. Proc Natl Acad Sci U S A, 2016. 113(15): p. 4039–44.

212. Moore, A.S. and E.L. Holzbaur, Dynamic recruitment and activation of ALS-associated TBK1 with its target optineurin are required for efficient mitophagy. Proc Natl Acad Sci U S A, 2016. 113(24): p. E3349–58.

213. Weil, R., et al., Role of Optineurin in the Mitochondrial Dysfunction: Potential Implications in Neurodegenerative Diseases and Cancer. Front Immunol, 2018. 9: p. 1243.

214. White, J., et al., Multifaceted roles of TAX1BP1 in autophagy. Autophagy, 2023. 19(1): p. 44–53.

215. Gao, L., et al., ABIN1 protein cooperates with TAX1BP1 and A20 proteins to inhibit antiviral signaling. J Biol Chem, 2011. 286(42): p. 36592–602.

216. Benyamin, B., et al., Cross-ethnic meta-analysis identifies association of the GPX3-TNIP1 locus with amyotrophic lateral sclerosis. Nat Commun, 2017. 8(1): p. 611.

217. Restuadi, R., et al., Functional characterisation of the amyotrophic lateral sclerosis risk locus GPX3/TNIP1. Genome Med, 2022. 14(1): p. 7.

218. Zhou, J., et al., TBK1 phosphorylation activates LIR-dependent degradation of the inflammation repressor TNIP1. J Cell Biol, 2023. 222(2).

219. Rasmussen, N.L., et al., The inflammation repressor TNIP1/ABIN-1 is degraded by autophagy following TBK1 phosphorylation of its LIR. Autophagy, 2023. 19(10): p. 2819–2820.

220. Le Guerroue, F., et al., TNIP1 inhibits selective autophagy via bipartite interaction with LC3/GABARAP and TAX1BP1. Mol Cell, 2023. 83(6): p. 927–941 e8.

221. Wu, S., et al., Structural basis for TNIP1 binding to FIP200 during mitophagy. J Biol Chem, 2024. 300(8): p. 107605.

222. White, J., et al., Phosphorylation of the selective autophagy receptor TAX1BP1 by TBK1 and IKBKE/IKKi promotes ATG8-family protein-dependent clearance of MAVS aggregates. Autophagy, 2024: p. 1–18.

223. Svenning, S., et al., Plant NBR1 is a selective autophagy substrate and a functional hybrid of the mammalian autophagic adapters NBR1 and p62/SQSTM1. Autophagy, 2011. 7(9): p. 993–1010.

224. Rasmussen, N.L., et al., NBR1: The archetypal selective autophagy receptor. J Cell Biol, 2022. 221(11).

225. Cozzi, M. and V. Ferrari, Autophagy Dysfunction in ALS: from Transport to Protein Degradation. J Mol Neurosci, 2022. 72(7): p. 1456–1481.

226. Yang, S., et al., Mutation analysis and immunopathological studies of PFN1 in familial and sporadic amyotrophic lateral sclerosis. Neurobiol Aging, 2013. 34(9): p. 2235 e7-10.

227. Farg, M.A., et al., C9ORF72, implicated in amytrophic lateral sclerosis and frontotemporal dementia, regulates endosomal trafficking. Hum Mol Genet, 2014. 23(13): p. 3579–95.

228. Renaud, L., et al., Key role of UBQLN2 in pathogenesis of amyotrophic lateral sclerosis and frontotemporal dementia. Acta Neuropathol Commun, 2019. 7(1): p. 103.

229. Alexander, E.J., et al., Ubiquilin 2 modulates ALS/FTD-linked FUS-RNA complex dynamics and stress granule formation. Proc Natl Acad Sci U S A, 2018. 115(49): p. E11485–E11494.

230. Zheng, T., S.K.K. Galagedera, and C.A. Castaneda, Previously uncharacterized interactions between the folded and intrinsically disordered domains impart asymmetric effects on UBQLN2 phase separation. Protein Sci, 2021. 30(7): p. 1467–1481.

231. Osaka, M., et al., Evidence of a link between ubiquilin 2 and optineurin in amyotrophic lateral sclerosis. Hum Mol Genet, 2015. 24(6): p. 1617–29.

232. Kim, J.E., et al., P2RX7-MAPK1/2-SP1 axis inhibits MTOR independent HSPB1-mediated astroglial autophagy. Cell Death Dis, 2018. 9(5): p. 546.

233. Alberti, S., et al., Granulostasis: Protein Quality Control of RNP Granules. Front Mol Neurosci, 2017. 10: p. 84.

234. Muranova, L.K., V.M. Shatov, and N.B. Gusev, Role of Small Heat Shock Proteins in the Remodeling of Actin Microfilaments. Biochemistry (Mosc), 2022. 87(8): p. 800–811.

235. Fox, L.M., et al., Huntington’s Disease Pathogenesis Is Modified In Vivo by Alfy/Wdfy3 and Selective Macroautophagy. Neuron, 2020. 105(5): p. 813–821 e6.

236. Clausen, T.H., et al., p62/SQSTM1 and ALFY interact to facilitate the formation of p62 bodies/ALIS and their degradation by autophagy. Autophagy, 2010. 6(3): p. 330–44.

237. Filimonenko, M., et al., The selective macroautophagic degradation of aggregated proteins requires the PI3P-binding protein Alfy. Mol Cell, 2010. 38(2): p. 265–79.

238. Popelka, H. and D.J. Klionsky, The emerging significance of Vac8, a multi-purpose armadillo-repeat protein in yeast. Autophagy, 2024: p. 1–4.

239. Park, J., et al., SIRT5-mediated lysine desuccinylation impacts diverse metabolic pathways. Mol Cell, 2013. 50(6): p. 919–30.

240. Kim, K.R., et al., S-Nitrosylation of cathepsin B affects autophagic flux and accumulation of protein aggregates in neurodegenerative disorders. Cell Death Differ, 2022. 29(11): p. 2137–2150.

241. Katunuma, N., Posttranslational processing and modification of cathepsins and cystatins. J Signal Transduct, 2010. 2010: p. 375345.

242. Xie, Z., et al., Cathepsin B in programmed cell death machinery: mechanisms of execution and regulatory pathways. Cell Death Dis, 2023. 14(4): p. 255.

243. Sen, N.E., et al., Search for SCA2 blood RNA biomarkers highlights Ataxin-2 as strong modifier of the mitochondrial factor PINK1 levels. Neurobiol Dis, 2016. 96: p. 115–126.

244. Singh, A., et al., Antagonistic roles for Ataxin-2 structured and disordered domains in RNP condensation. Elife, 2021. 10.

245. Shibata, H., D.P. Huynh, and S.M. Pulst, A novel protein with RNA-binding motifs interacts with ataxin-2. Hum Mol Genet, 2000. 9(9): p. 1303–13.

246. Ponnambalam, S. and S.A. Baldwin, Constitutive protein secretion from the trans-Golgi network to the plasma membrane. Mol Membr Biol, 2003. 20(2): p. 129–39.

247. Yin, D.M., et al., Both the establishment and maintenance of neuronal polarity require the activity of protein kinase D in the Golgi apparatus. J Neurosci, 2008. 28(35): p. 8832–43.

248. Bisbal, M., et al., Protein kinase d regulates trafficking of dendritic membrane proteins in developing neurons. J Neurosci, 2008. 28(37): p. 9297–308.

249. Czondor, K., et al., Protein kinase D controls the integrity of Golgi apparatus and the maintenance of dendritic arborization in hippocampal neurons. Mol Biol Cell, 2009. 20(7): p. 2108–20.

250. Fujishima, K., et al., Principles of branch dynamics governing shape characteristics of cerebellar Purkinje cell dendrites. Development, 2012. 139(18): p. 3442–55.

251. Bencsik, N., et al., Protein kinase D promotes plasticity-induced F-actin stabilization in dendritic spines and regulates memory formation. J Cell Biol, 2015. 210(5): p. 771–83.

252. Oueslati Morales, C.O., et al., Protein kinase D promotes activity-dependent AMPA receptor endocytosis in hippocampal neurons. Traffic, 2021. 22(12): p. 454–470.

253. Besirli, C.G. and E.M. Johnson, Jr., The activation loop phosphorylation of protein kinase D is an early marker of neuronal DNA damage. J Neurochem, 2006. 99(1): p. 218–25.

254. Krueger, D.D., E.K. Osterweil, and M.F. Bear, Activation of mGluR5 induces rapid and long-lasting protein kinase D phosphorylation in hippocampal neurons. J Mol Neurosci, 2010. 42(1): p. 1–8.

255. Liliom, H., et al., Protein kinase D exerts neuroprotective functions during oxidative stress via nuclear factor kappa B-independent signaling pathways. J Neurochem, 2017. 142(6): p. 948–961.

256. Pose-Utrilla, J., et al., Excitotoxic inactivation of constitutive oxidative stress detoxification pathway in neurons can be rescued by PKD1. Nat Commun, 2017. 8(1): p. 2275.

257. Johnson, J.R. and J.W. Barclay, C. elegans dkf-1 (Protein Kinase D1) mutants have age-dependent defects in locomotion and neuromuscular transmission. MicroPubl Biol, 2023. 2023.

258. Uesugi, A., et al., Involvement of protein kinase D in uridine diphosphate-induced microglial macropinocytosis and phagocytosis. Glia, 2012. 60(7): p. 1094–105.

259. Yang, Y., et al., TDP-43 levels in the brain tissue of ALS cases with and without C9ORF72 or ATXN2 gene expansions. Neurology, 2019. 93(19): p. e1748–e1755.

260. Goedert, M., Tau protein and the neurofibrillary pathology of Alzheimer’s disease. Trends Neurosci, 1993. 16(11): p. 460–5.

261. Lee, G., Tau and src family tyrosine kinases. Biochim Biophys Acta, 2005. 1739(2-3): p. 323–30.

262. Sieben, A., et al., The genetics and neuropathology of frontotemporal lobar degeneration. Acta Neuropathol, 2012. 124(3): p. 353–72.

263. Neumann, M., et al., Ubiquitinated TDP-43 in frontotemporal lobar degeneration and amyotrophic lateral sclerosis. Science, 2006. 314(5796): p. 130-3.

264. Li, H.Y., et al., Hyperphosphorylation as a defense mechanism to reduce TDP-43 aggregation. PLoS One, 2011. 6(8): p. e23075.

265. Wang, A., et al., A single N-terminal phosphomimic disrupts TDP-43 polymerization, phase separation, and RNA splicing. EMBO J, 2018. 37(5).

266. Gruijs da Silva, L.A., et al., Disease-linked TDP-43 hyperphosphorylation suppresses TDP-43 condensation and aggregation. EMBO J, 2022. 41(8): p. e108443.

267. Haider, R., et al., Phosphomimetic substitutions in TDP-43’s transiently alpha-helical region suppress phase separation. Biophys J, 2024. 123(3): p. 361–373.

268. Kim, K.Y., et al., A phosphomimetic mutant TDP-43 (S409/410E) induces Drosha instability and cytotoxicity in Neuro 2A cells. Biochem Biophys Res Commun, 2015. 464(1): p. 236–43.

269. Paron, F., et al., Unraveling the toxic effects mediated by the neurodegenerative disease-associated S375G mutation of TDP-43 and its S375E phosphomimetic variant. J Biol Chem, 2022. 298(8): p. 102252.

270. Hofweber, M. and D. Dormann, Friend or foe-Post-translational modifications as regulators of phase separation and RNP granule dynamics. J Biol Chem, 2019. 294(18): p. 7137–7150.

271. Ainani, H., et al., Liquid-liquid phase separation of protein tau: An emerging process in Alzheimer’s disease pathogenesis. Neurobiol Dis, 2023. 178: p. 106011.

272. Truett, G.E., et al., Preparation of PCR-quality mouse genomic DNA with hot sodium hydroxide and tris (HotSHOT). Biotechniques, 2000. 29(1): p. 52, 54.

273. Eng, J.K., T.A. Jahan, and M.R. Hoopmann, Comet: an open-source MS/MS sequence database search tool. Proteomics, 2013. 13(1): p. 22–4.

274. Huttlin, E.L., et al., A tissue-specific atlas of mouse protein phosphorylation and expression. Cell, 2010. 143(7): p. 1174–89.

275. Villen, J., et al., Large-scale phosphorylation analysis of mouse liver. Proc Natl Acad Sci U S A, 2007. 104(5): p. 1488–93.

276. Livak, K.J. and T.D. Schmittgen, Analysis of relative gene expression data using real-time quantitative PCR and the 2(-Delta Delta C(T)) Method. Methods, 2001. 25(4): p. 402–8.

277. Klann, K., G. Tascher, and C. Munch, Functional Translatome Proteomics Reveal Converging and Dose-Dependent Regulation by mTORC1 and eIF2alpha. Mol Cell, 2020. 77(4): p. 913–925 e4.

278. Sutandy, F.X.R., et al., A cytosolic surveillance mechanism activates the mitochondrial UPR. Nature, 2023. 618(7966): p. 849–854.

279. Szklarczyk, D., et al., STRING v11: protein-protein association networks with increased coverage, supporting functional discovery in genome-wide experimental datasets. Nucleic Acids Res, 2019. 47(D1): p. D607–d613.

